# CONFESS: Fluorescence-based single-cell ordering in R

**DOI:** 10.1101/407932

**Authors:** Efthymios Motakis, Diana H.P. Low

## Abstract

Modern high-throughput single-cell technologies facilitate the efficient processing of hundreds of individual cells to comprehensively study their morphological and genomic heterogeneity. Fluidigm’s C1 Auto Prep system isolates fluorescence-stained cells into specially designed capture sites, generates high-resolution image data and prepares the associated cDNA libraries for mRNA sequencing. Current statistical methods focus on the analysis of the gene expression profiles and ignore the important information carried by the images. Here we propose a new direction for single-cell data analysis and develop CONFESS, a customized cell detection and fluorescence signal estimation model for images coming from the Fluidigm C1 system. Applied to a set of HeLa cells expressing fluorescence cell cycle reporters, the method predicted the progression state of hundreds of samples and enabled us to study the spatio-temporal dynamics of the HeLa cell cycle. The output can be easily integrated with the associated single-cell RNA-seq expression profiles for deeper understanding of a given biological system. **CONFESS** R package is available at Bioconductor (http://bioconductor.org/packages/release/bioc/html/CONFESS.html).

## 1 Introduction

Over the last few years there have been considerable developments towards the elucidation of the progression dynamics of complex molecular mechanisms such as cell cycle and proliferation. The recently developed single-cell high-throughput technologies have facilitated the time and cost effective analysis of large cell populations and have led to profound biological discoveries setting high expectations in the research of human diseases. Notably, Fluidigm’s C1 microfluidics system attempts to automatically isolate each cell into an individual reaction chamber (capture site) located within an integrated fluidic circuit (IFC) or chip. Typically, the chips are designed with either 96 or 800 chambers. The chambers IDs of the former design are derived from a combination of A-H and 1-12 characters. The captured cells are stained and examined for viability before subjected to downstream analysis. Due to technological noise or other errors, there is a non-negligible probability that some chambers might end up being empty or contain more than one cell either on the site or in the vicinity (doublets).

Essentially, the raw data are noisy images of the irregular 2-dimensional characteristics of the stained cells in one or more channels. The images carry important experimental information on the data quality and the cell properties that impacts the accuracy of the downstream predictions (cell phenotypes). The first analytic step calls for cell detection and fluorescence signal estimation in the capture site. Thermo Scientific Cellomics HCS Solution carries out these tasks via a commercial image analysis platform. Alternatively, one could refer to a wide range of open-source CRAN packages (https://cran.r-project.org) for general image processing such as **adimpro** [1], **EBImage** [2] and **raster** [3]. Since none of these choices are designed for the analysis of the Fluidigm C1 data, one has to be cautious of estimation errors (false positives / negatives) and develop additional procedures for quality control and recovery of missing information [4]. In a typical single-cell high-throughput study, the captured cells are lysed, their RNA is reverse transcribed and massive parallel sequencing interrogates their transcriptional characteristics. Recent works have shown that most transcripts exhibit high cell-to-cell variability [5–7] and suggests that relatively large sample sizes (typically 200 cells or more) are often required to make accurate conclusions [8,9]. Thus, cell fluorescence signals coming from multiple 96-chamber chips, often processed in runs by different experimenters across several days and machines, should be appropriately integrated to minimize unwanted non-biological effects.

A challenging task that have been extensively addressed in single-cell studies is the reconstruction of a given biological mechanism through the analysis of the pseudo-dynamic system of cell progression. Modern statistical methods including **TSCAN** [10], **Monocle** [11], **Oscope** [12], **Waterfall** [13], **SCUBA** [14], **DeLorean** [15], **SLICE** [16] and **SCENT** [17] model directly the single-cell RNA-seq gene expression profiles to provide a pseudo-temporal ordering of the cells and subsequently study significant changes on their transcriptome. None of these approaches exploits the image information except for quality control purposes [6,18]. Exceptions are the recently developed **Cycler** [19] and **Wanderlust** [20] that estimate cell cycle progression within a multi-dimensional feature space. **Cycler** utilizes the image data of the nucleus of fixed cells combined with other single-cell measurements. **Wanderlust** performs trajectory detection (but not clustering) via graph-based ensembles using explicitly mass or flow cytometry generated data. Although quantitatively richer, the quality control of such datasets is a difficult task.

Models that analyse flow-cytometry (FC) fluorescence data such as **SamSPECTRAL** [21], **flowMerge** [22], **flowMeans** [23] and **flowPeaks** [24] apply sophisticated 2-dimensional clustering techniques to predict homogeneous cell groups from the signals. By themselves, these models do not generate enough information to reconstruct a dynamic biological mechanism and enable deeper transcriptomic analysis. In this work, we address the above issues in our novel statistical model whose multiple steps are integrated into the *C*ell *O*rderi*N*g (by) *F*luor*ES*cence *S*ignal **CONFESS** R package. **CONFESS** performs image analysis and fluorescence signal estimation for data coming from the emerging Fluidigm C1 technology. It collects extensive information on the cell morphology, location and signal that can be used for quality control and phenotype prediction. If applicable, it normalizes and uses the signals for unsupervised cell ordering (pseudotime estimation) and 2-dimensional clustering via scalar projection, change-point analysis and Data Driven Haar Fisz transformation for multivariate data (*DDHFmv*). Its output can be easily integrated with available single-cell RNA-seq (or other) expression profiles for the subsequent study. **CONFESS** is suitable for all single-cell applications utilizing living cells expressing a set of appropriate fluorescence reporters that can determine their biological state [25]. Here, it is fully implemented to study the HeLa cell cycle using a modified version of the *F* luorescent, *U* biquitination-based *C* ell *C* ycle *I* ndicator (Fucci) transgenes system [26] (developed on mouse embryonic stem cells) where the order of the red and green color transitions is similar to the original Fucci [25].

This paper is organized as follows: Section 2 gives a detailed description of **CONFESS** and its distinct steps, i.e. the fluorescence estimation and quality control (Section 2.1), the background and run effect correction (Sections 2.2 - 2.3) and the spatio-temporal data analysis of a dynamic biological process (Section 2.4). Section 3 illustrates its functionality on real and simulated data. Section 4 shows the robustness of our predictions by cross-validation that allows assessing potential outliers. Section 5 compares our methodology with existing flow cytometry based clustering approaches only since, at this moment, the authors do not have permission to publish the singe-cell RNA-seq data of the HeLa experiment. Finally, Section 6 highlights the main conclusions of this work.

## 2 Mathematical formulation of CONFESS

This section describes **CONFESS** and its three analytical steps. Briefly, the first step utilizes image analysis functions from **EBIimage** and **raster** R packages to locate the cell on the C1 chip, estimate its raw fluorescence signal and perform quality control. If the signal is too weak to detect, the model applies pattern recognition on the architecture of the C1 chip to locate the capture site and measure the signal within. The second step performs background and, if needed, run effect signal adjustment by finite mixture models [27] in order to minimize systematic, non-biological sources of variation such as chip-to-chip and image-specific errors [28]. The third step performs pseudo-temporal cell ordering via scalar projection and change-point analysis on appropriately transformed signals to study the dynamics of a biological mechanism such as the cell cycle.

### 2.1 Identification of cell location and morphology, raw signal estimation and quality control

We describe **CONFESS** in terms of our HeLa dataset [28] where the cells express the red fluorescence signal in the early G1, G1 (growth) and S (synthesis) phases. The green fluorescence is then activated at S and remains at high levels until G2 and M (replication). At the end of cell division both signals reach their minimum levels. Each potential single-cell sample is represented by a C1 image triplet *i, i* = 1,…,*I* (*I* = 378), generated from Cellomics ArrayScan VTI High Content Analysis Reader. The three types of C1 images are: (i) the Bright Field (*BF*) image, *i*^*BF*^, (ii) the Green, *i*^*Green*^, channel and (iii) the Red, *i*^*Red*^, channel. The BF images contain the C1 chip information (chip pattern and the location of the capture site) while the channel images show the morphology and the signal of the stained cells relative to the background. For simplicity, we will use the term ‘image’ instead of ‘image triplet’ and refer to channel-specific images as *i*^*c*^, *c* ∈ {*Green, Red*}, when necessary.

### 2.1.1 The input images or converted data matrices

**CONFESS** expects as input the pixel signals of the raw image data in a matrix form. The user can convert the images into raw signal matrices (BMP/JPEG/PNG to .txt) directly, or independently by any standard tool such as ImageJ (http://fiji.sc/). In this project, we analyzed high-resolution .C01 format images (converted into .txt by ImageJ) of dimension 512 × 512 each. The signals of each Green and Red channel matrix are independently normalized to range within [0,1] as:

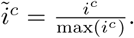

The elements of each 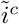 are binarized by the indicator function

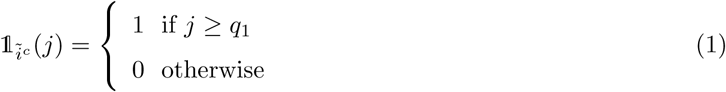

for every element *j* of 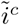 and a series of predefined cut-offs 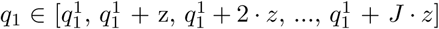 where *z* is a small constant (e.g. *z* = 0.02), 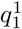 is a starting value (e.g. 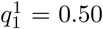) and *J* an arbitrary value such that 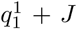. *z* ≤ 1.

### 2.1.2 The standard spot identification functions

In the following paragraphs we discuss how spot identification leads to cell estimation through quality control. The term *spot* is used to highlight the uncertainty of **CONFESS**’s first-step estimates that could correspond either to an artificial cell-like object or a true cell.

Denoted as 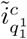, the binary signal matrix of channel *c* is obtained by the minimum cut-off 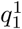 in (1). On this data, **CONFESS** identifies the area of a single bright spot (that may correspond to the cell of interest) of any size, shape and at any location on the image. If 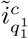 does not succeed the algorithm iterates on a new data matrix 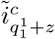. The process stops at the first cut-off that meets the criterion.

The algorithm will not necessarily identify a single spot per image *i*^*c*^. In fact, in noisy data or images with artefacts, **CONFESS** could potentially return zero spots (e.g. when the signal is too weak or the noise is too high) or store the coordinates and the area sizes of several spots (e.g. when artificial spot-like regions appear). In our data, such cases occur in approximately 20% of the samples. Thus, we seek ways to combine the above spot measures across channels into a single set of characteristics reflecting the location, the size and the signal of a potentially true cell.

#### 2.1.3 First-step spot identification via consensus estimation and BF image modelling

This paragraph describes how **CONFESS** obtains the *first-step spot estimates*. The supporting figures, reflecting possible estimated scenarios, are based on simulated data from a *Poisson* 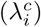 + *Normal*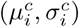 model [29]. The simple scenarios can be resolved by consensus estimation, i.e. the spot locations from the two channels are combined to a consensus output. In complex cases of multiple spots we apply BF image modelling to find patterns on the C1 chip architecture, locate the capture site and retrieve the estimates of the closest spot in it (if it exists). These first-step estimates will be subjected to quality control to ensure that the identified spot represents a true cell.

##### Maximum one spot in each channel

First we discuss Case 1A of Figure 1 where **CONFESS** locates exactly one spot in each channel *c* of image *i* such that:

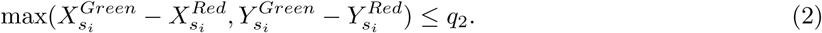

**Figure 1:**
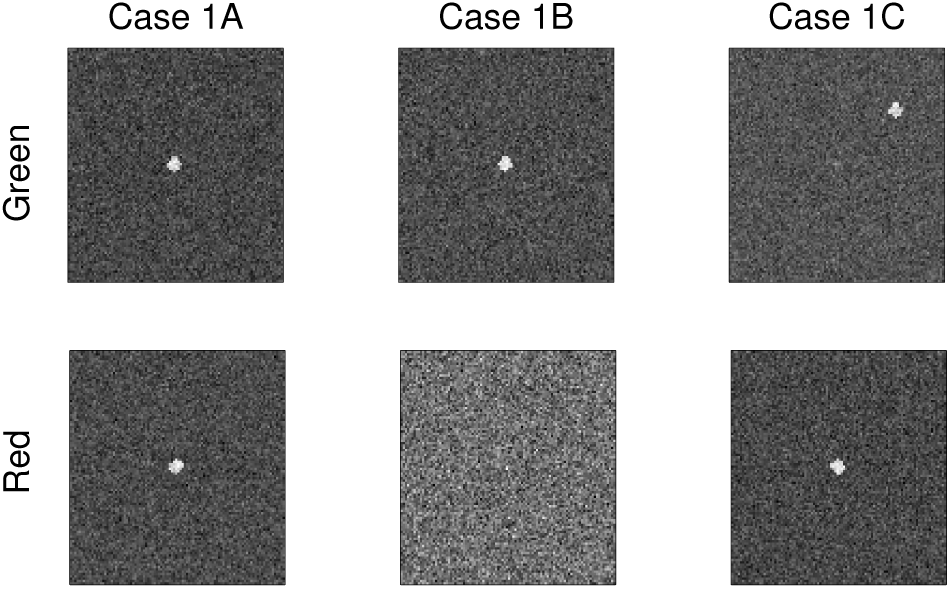
Simulated examples of two-channel images with maximum one spot per channel. Case 1A: The spots are matched suggesting the existence of a cell. Case 1B: The spot appears only in the Green channel. Case 1C: The channels contain unmatched spots and at least one is false a positive.

Here,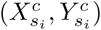 is the 2-dimensional coordinates of the spot’s *s* brightest pixel on *i* and *c, s* = 1, …, *S* to allow for multiple spots per image and *q*_2_ is a user-specified cut-off (in our analysis *q*_2_=7). The image (and the associated signal) data can be easily simulated by simcells() whose detailed description is provided in Section 3.

If (2) is satisfied, implying that the spots are close to each other across channels (a matching pair), **CONFESS** will return the following set of estimates:

1. *Spot center*: This is a representative spot location on the image and it is estimated as:

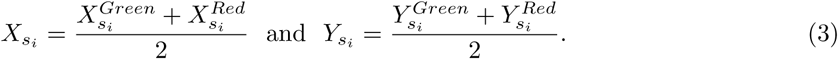 Note that *q*_2_ is small enough to ensure that the center will not be estimated on a background region;
2. *Spot size*: Let 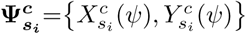 *ψ* = 1, &, Ψ, be the set of 2-dimensional pixel coordinates of 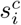. The total spot area is the union

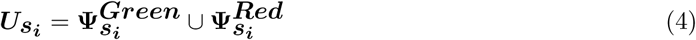

and the spot size is the cardinality of the union set, i.e. ——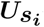 ——. There are no assumptions on the spot shape;
3. *Cell signal*: The spot signal (foreground) is estimated as the average of the pixel signals 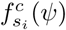

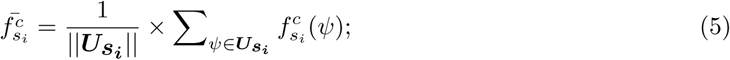
4. *Background signal*: The background signal serves as an image-specific normalization metric. It is estimated as the average signal of *B* (default *B*=100) pseudo-spots of the same shape and size at random locations on the image *i* of channel *c*. Mathematically the background signal is:

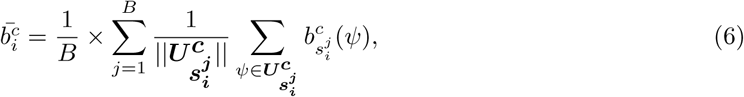

where 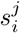 is the *j*th pseudo-spot of *i*, 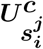 is the set of pseudo-spot’s *j* pixel coordinates on *i* and *c* such that 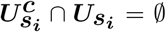 and 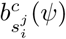 is the signal of (background) pixel *ψ* of pseudo-spot 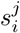 on *c*. Notice that (6) summarizes the pseudo-spot signals into a common background signal;
5. *Signal-to-Noise ratio*: The signal-to-noise ratio evaluates the difference between 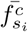 and 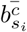 as:

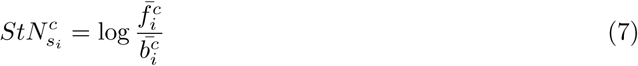

and eventually assesses the existence, the viability and the biological state of a cell. For example, dead cells are expected not to reject the hypothesis 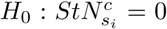 at *α* = 5% by the Wilcoxon test (thus the foreground is statistically equal to the background).

Case 1B of Figure 1 illustrates the case of a single spot in only one channel. Were that spot a true cell, this would suggest its cell cycle phase. Here, **CONFESS** finds the spot coordinates (3) - (4) from the ‘occupied’ image and apply them on both channels to retrieve the estimates (5) - (7).

In Case 1C of Figure 1 there is exactly one spot per image and (2) is not met. This information is not sufficient to link a spot to a cell since *at least one* of the two spots is a false positive. Such cases are resolved by BF image modelling that identifies the chamber’s location and matches it to the coordinates of the closest spot that satisfies (2). Then the algorithm processes the information as in Case 1B. If unsuccessful, the estimation is solely based on BF image modelling.

##### Only one multi-spot channel

The simplest scenario is illustrated in Case 2A of Figure 2 where (2) is satisfied for one pair (the spots on the center) that will be linked to a potential cell. **CONFESS** will process the pair as in Case 1A. The uncertain cases of Case 2B and 2C (where (2) is not met) are resolved as in Case 1C using a combination of BF image modelling and identification of the chamber’s location.

**Figure 2:**
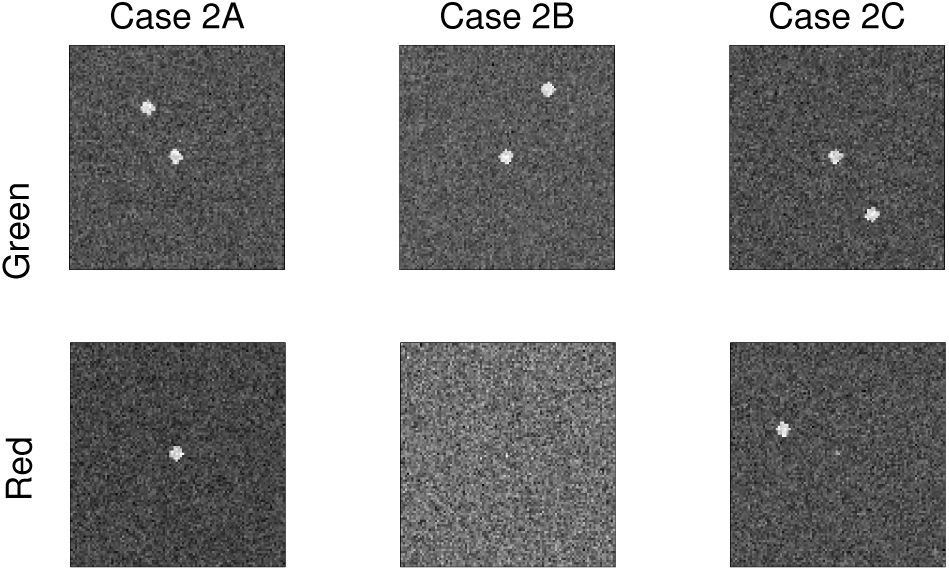
Simulated examples of two-channel images only one of which contains more than one spot. Case 2A: The matched spot pair suggests the existence of a cell on the image center. The other spot (Green channel) can be considered as noise. Case 2B: Two spots appear on the Green channel but it is unclear which one is associated to a cell. Case 2C: The channels contain unmatched spots and at least one is a false positive.

##### Two multi-spot channels

The simplest scenario is shown in Case 3A of Figure 3 with one pair satisfying (2) (the spots on the center) that enables **CONFESS** to use the methodology of Case 1A. Due to image scanning or other technology errors, it is possible that (2) is satisfied by two spot pairs simultaneously as in Case 3B. This may occur either because two cells were captured in the chip or due to a technical artefact on the image that generated a cell-like spot. The algorithm will give priority to the pair with the largest |***Us*_*i*_** | (to eliminate bright spot-like speckles) but it will also keep the estimates of the second spot as an alternative solution. Note that in the presence of two cells, the image will be flagged as contaminated and will be discarded. Finally, Case 3C shows an extreme example of no matched coordinates for multiple spots which is addressed as in Case 1C with BF modelling.

**Figure 3:**
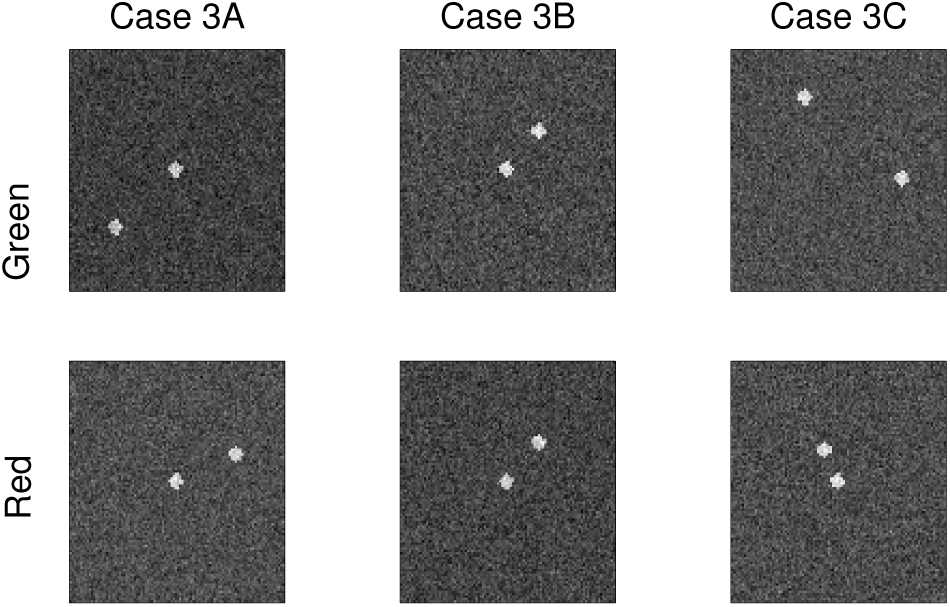
Simulated examples of two-channel multi-spot images. Case 3A: The matched spot pair suggests the existence of a cell on the image center. The other spots can be considered as noise. Case 3B: The existence of two matching spot pairs suggests image contamination by a cell-like artefact or a chip failure to capture only one cell. Case 3C: The channels contain unmatched spots and at least one is a false positive.

##### Spotless images in both channels

In the case of empty images **CONFESS** will directly apply BF image modelling. Figure 4 shows the architecture of two C1 chips with their characteristic capture sites (a small curved slot). The location of a capture site varies according to:

1. The chamber ID: chamber IDs of the form x 2/5/7/9/10/12 are facing right (Figure 4a) while the x 1/3/4/6/8/11 ones are facing left (Figure 4b). Here ‘x’ denotes one of the letters A-H of the original chamber ID;
2. The frame: even for the cases of similar chamber IDs the chamber is roughly located at the same coordinates due to image frame shifts.

**Figure 4:**
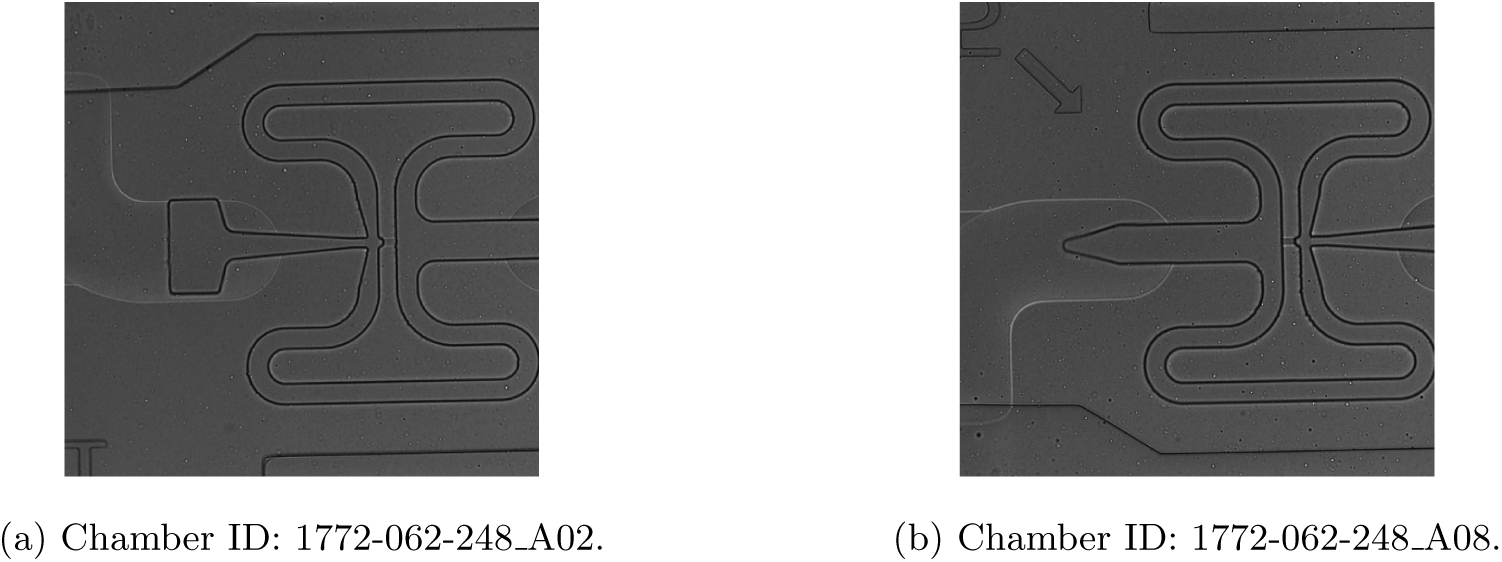
An example of the architecture of Fluidigm’s C1 chip. The directionality of the capture site depends on the chamber ID.

Nevertheless, most of the chip’s features such as the dark straight-like horizontal and vertical lines will always be present. BF modelling locates these lines by a run-based algorithm:

1. Produce the [0,1]-normalized version of the BF matrix as:

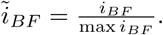 The pixels with features of interest (dark lines) tend to have low normalized signals. Consider a series of small *q*_3_ cutoffs, typically *q*_3_ = [0.3, 0.32, 0.34, …,0.5]. Apply the first one,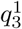, to (1) to set the darkest features of 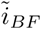 to zero;
2. Find runs of 0’s in rows and columns and store their length. The rows and columns with the maximum run length correspond to features. Since on the C1 chip the distances among the straight lines are standard, the algorithm will find the row and column sets that satisfy these distances (fed in the algorithm by default);
3. If 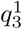 converges to a unique set of rows and columns that agree with the C1 chip specifications, **CONFESS** will stop the process and simply estimate the chamber’s location by integrating the information. Otherwise, it will iteratively select the next cut-off until convergence;
4. Consider a user-defined rectangular region (default is 7 × 7) and perform foreground estimation (5) for its pixels. Consequently, the spot size is always set to a user-defined constant (here 7^2^ = 49). The background and signal-to-noise ratio are defined as before.

To sum up, in the presence of a spot pair that satisfies (2), the estimates (3) - (7) are exclusively fluorescence based. Otherwise, BF image modelling will locate the chamber and apply (3) - (7) accordingly. These estimates will assist **CONFESS** to determine the existence of a spot that may correspond to a viable cell. The success of the pipeline depends on the image quality. The results and the simulations (Section 3) indicate the high accuracy of our approach.

#### 2.1.4 From spot to the cell raw signal by quality control and re-estimation

The basic assumption of the above model is that the fluorescence signals, sometimes integrated with BF modelling, tend to point to true cells. Still, the estimation will sometimes fail by giving unusual (*X*_*s*_*i, Y*_*s*_*i*) coordinates. Such a hypothetical scenario can occur if, for example, in Case 2A the true cell is represented by the single, unmatched spot (top). In order to avoid such errors, **CONFESS** incorporates the following quality control process:

1. For each of the runs and chamber sets (x 1/3/4/6/8/11 and x 2/5/7/9/10/12), plot all *I*_*run,chambID*_-specific spot center locations (Figure 5);

**Figure 5:**
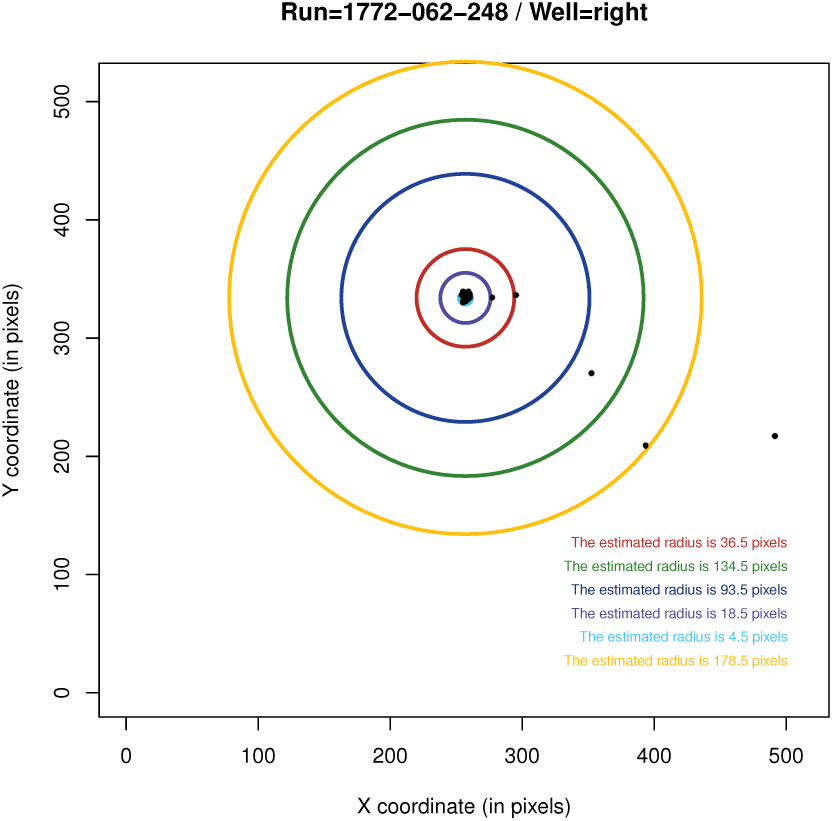
Graphical visualization of the estimated spot locations (*X*_*s*_*i, Y*_*s*_*i*) for run ID 1772-062-248 and chamber IDs facing right (x 2/5/7/9/10/12). The concentric circles are generated for outlier detection. Similar plots are produced for each runs and chamber directionality.
2. Assuming that the majority of the locations are well-estimated (the bulk of points), estimate the 2-dimensional median *X*_*cent*_, *Y*_*cent*_ (the centroid shown by the red cross) and calculate the Euclidean distance between each point and the centroid as:

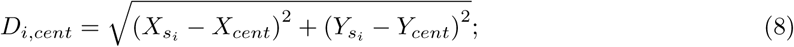
3. Apply the Grubbs test for outliers on *D*_*i,cent*_ to find unusual spot locations at various significant levels. The details and options are given in Section 3;
4. The flagged images that have been originally estimated by fluorescence signal are re-estimated by BF modelling. An exception holds for Case 3B where the second spot estimates substitute the original ones. The flagged images that have been originally estimated by BF modelling are either outliers and are not processed further or are subjected to *forced adjustment* that sets the chamber’s central location at (*Xcent, Ycent*).

The above procedure gives the final locations and raw cell fluorescence estimates that are simply denoted by 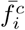 and 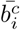 (the spot subscript is dropped). The number and locations of all secondary spots (potentially false positives) are data quality indicators and are also stored. We suggest that all **CONFESS** figures should be manually inspected for verification.

#### 2.1.5 Output and further notation

The output is cell-specific and consists of several quantities describing mainly the location (cell center (*X*_*i*_, *Y*_*i*_) and cell size |***U*** |) and the fluorescence characteristics (foreground 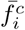 and background 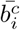 signals) of each cell. Additional quality control measures (signal-to-noise ratio 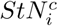 with its associated significance metrics, a quality control index highlighting potential outliers and the coordinates of all secondary spots) are also included. Next, we will only discuss the estimates of a single cell on each image *i* and thus we will use *i* = 1, *…, I* to denote both the images and the cells.

### 2.2 Background adjustment of the raw fluorescence signals

The background corrected cell signal, 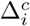, is a summarized version of the raw foreground 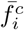 and background 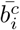 signals. The simplest approach to obtain it is through the *subtraction model*, i.e. 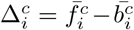, that generates undesirable zero or negative values when 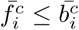 and does not correct for the ambient intensity surrounding the cell itself [30,31]. An alternative statistical approach for background correction is the *normexp* model that was developed by Ritchie et al [32] and computationally refined by Silver et al [33]. Briefly, *normexp* fits the background subtracted signals 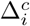 in a convolution model of two random variables: one exponentially distributed representing the true signal 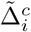 and the other normally distributed representing the background noise:

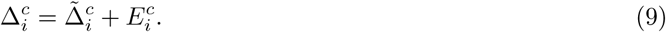

The estimation of the background corrected signal 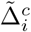 is performed by Maximum Likelihood using saddle-points [33]. The above model is essentially a smooth monotonic transformation of the background subtracted intensities such that all the corrected intensities are positive.

### 2.3 The integrated run effect and background adjustment model

By itself, model (9) cannot correct systematic chip-to-chip (run) variations. For this reason, in our previous work [28] we developed a two-step approach that first adjusts for run effects on multi-modal 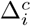 signals (Figure 6) and then uses *normexp* to obtain the run and background adjusted 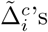.

**Figure 6:**
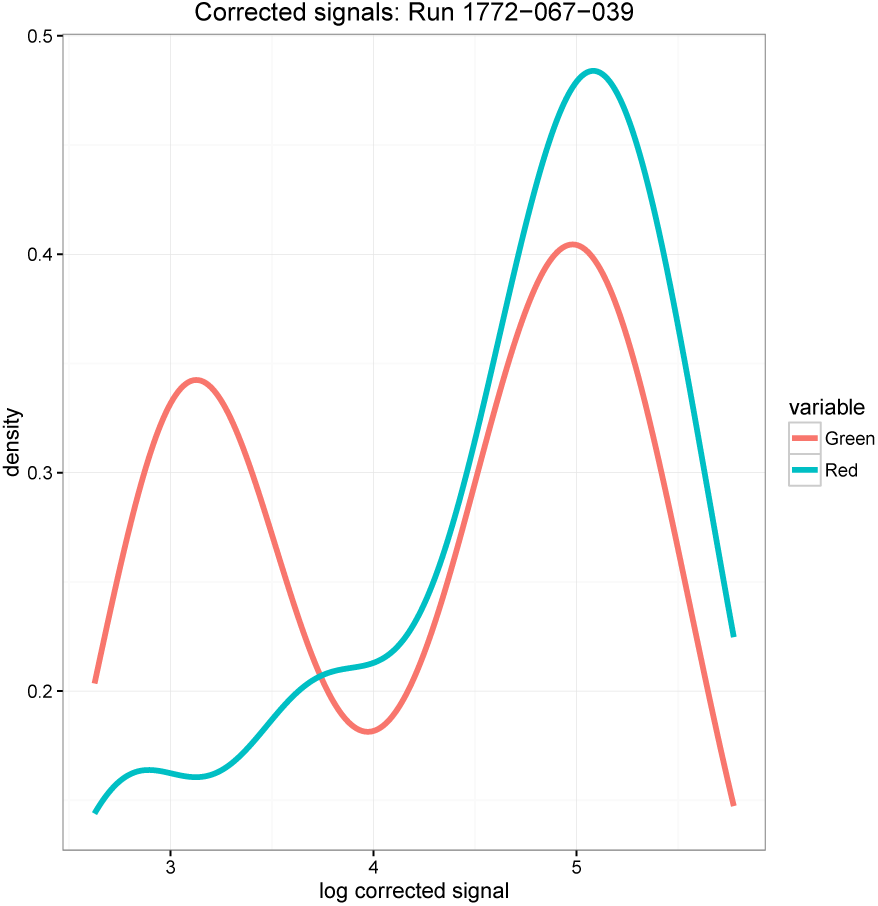
The densities of the background corrected fluorescence signals of run 1772-067-039.

The model utilizes a latent class Bayesian finite mixtures of regression model of the form:

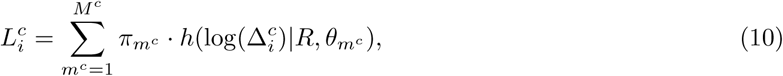

where *M*^*c*^ is the unknown number of mixtures to be estimated in each channel *c* from the 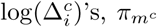 is the prior probability of the channel-specific component *m*^*c*^, 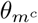 is a component-specific parameter vector to be estimated and *h*(.) is the conditional density of log(Δ^*c*^) on the runs *R* and 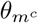. Model (10) with constraints:

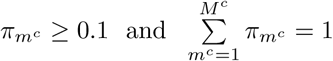

was originally described in Leisch et al [27] and it is currently applied in **flexmix**. Setting 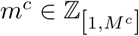 allows each channel to have different number of mixtures. Model (10) is fitted for a number of *M*^*c*^ components (e.g. we set *m*^*c*^ ∈ 𝕫 _[1,3]_ after visual inspection) and carries out model selection to estimate the optimal *m*^*c*^ using the Bayesian Information Criterion, estimated by the Expectation-Maximization algorithm repeatedly 50 times using different starting values [27]. This procedure predicts the categorical variable *L*^*c*^ of the number of mixtures and fits it into a multi-factorial ANOVA with interactions:

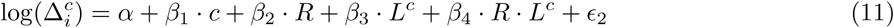

that adjusts for the chip-to-chip variation by removing the main effect of the runs, *β*_2_, and the interaction effect between the runs and each component, *β*_4_, as:

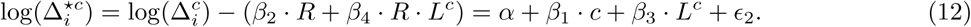

For data coming from a single run, the reduced right hand side version of (12) can be directly fitted. In the case of 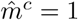 in both channels, model (11) becomes a 2-way ANOVA:

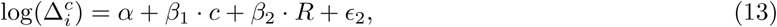

where only *β*_2_ effect is removed. Replacing the left hand side of (9) with log 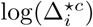 of (12) and fitting the model obtains the adjusted fluorescence signals 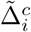.

### 2.4 Fluorescence signals model for cell ordering and clustering

**CONFESS** initiates cell ordering by sorting the 2-dimensional 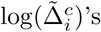 according to a predefined or predicted path. The path can be inferred from the characteristics of the experimental technology, e.g. the properties of the fluorescence reporters [25], the associated biological study and the spatio-temporal behaviour of the estimated signals. Figure 7 shows an example coming from the analysis of the data.

**Figure 7:**
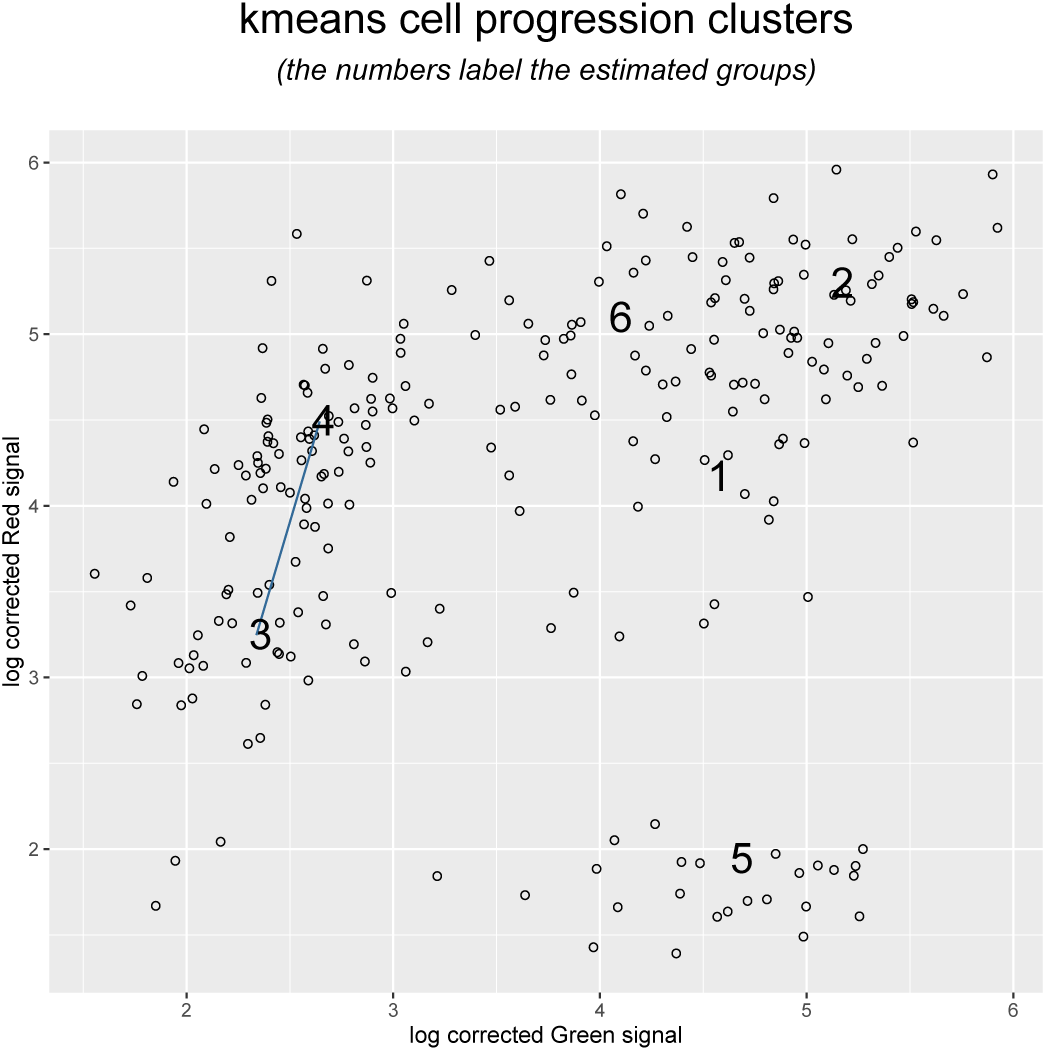
Clustering the corrected channel signals by 2-dimensional *k*-means to obtain an initial estimation of cell progression clusters. Parameter *k* is estimated by the gap statistic. The numbers are placed at the cluster centroids. The original plot by **CONFESS** does not include the straight line, which is added here for illustration purposes (see Section 3.1; *Unsupervised cell ordering and grouping*).

The 2-dimensional scatterplot of the log-transformed adjusted signals confirms the previously described clockwise path of cell cycle progression [25]. First, we use 2-dimensional *k*-means clustering, with *k* estimated by the gap-statistic [34], to obtain the initial clusters, denoted by *k* = 1, …, *K* (here *K* = 5). Then, we estimate the *K* 2-dimensional medians (numbers in Figure 7) as:

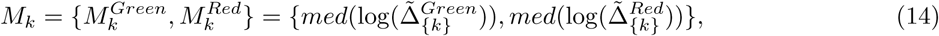

where 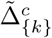 is the set of adjusted cell signals of the k cluster in c. Visually, the *M*_*k*_’s are ordered into a cell cycle path that is computationally retrieved below (paragraph *Estimation of the progression path*). Starting from any initial group *k*, we can move either clockwise or anticlockwise to the next group *k* + 1 whose single cells *i* (denoted by *i*(*k* + 1)) are to be ordered by pseudotime. The pseudotime, *T*_*i*(*k*+1)_, is estimated by scalar projection.

We define 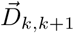 as the vector that connects *M*_*k*_ and *M*_*k*+1_ (the straight line in Figure 7), and 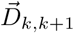 as the vector that connects the 2-dimensional log-signal of *i*(*k* + 1) and 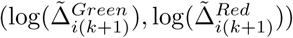, with *M*_*k*+1_. We compute the orthogonal projection of each *i*(*k* + 1) onto 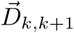 as:

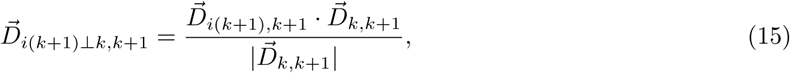

where

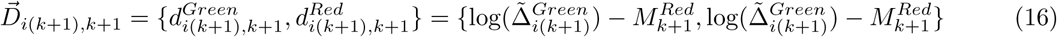

and

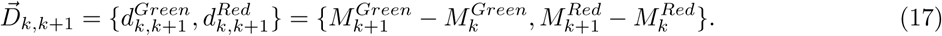

wrapping up (15) - (17) we obtain:

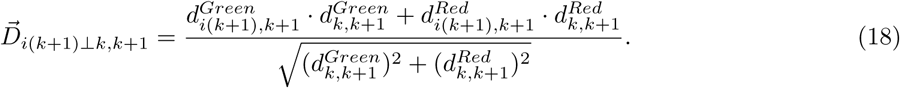

Following the progression path we can estimate the 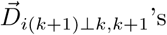 for all *i*(*k*+1) and all *k, k*+1 neighbouring clusters, *k* = 1,.., *K* - 1. In circular paths, to compute the orthogonal projection of the *i*(1)’s, i.e. the cells of the initial cluster, we consider the pair *K*, 1 and estimate (18) as 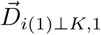 where *K* is the final cluster of the path.

The above framework implies that small 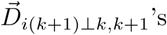 are located farther from *M*_*k*+1_ and closer to *M*_*k*_. To provide a more accurate representation of the dynamic changes across phases we developed an *inter-cluster smoother* that mimics the above in the opposite direction. For each *i*(*k* + 1) we estimate

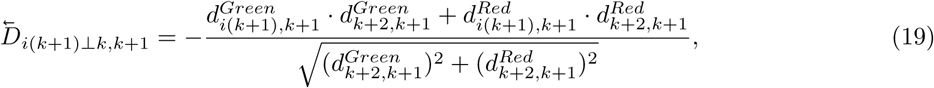

where the minus sign defines that (19) assigns small 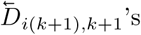 to *i*(*k* + 1)’s that are closer to *M*_*k*+2_ than to *M*_*k*+1_. The average of (18) - (19)

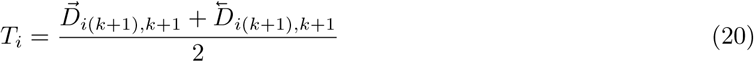

denotes the estimated cell ordering pseudotimes. **CONFESS**’s predictions can be fed to any standard parametric or non-parametric model for further analysis on gene expression, biomarker discovery via machine learning, motif activities, co-regulation networks and other biologically interesting tasks.

### 2.4.1 Estimation of the progression path

As noted above, the 2-dimensional centroids *M*_*k*_, *k* = 1, *…, K*, summarize the fluorescence changes across clusters and can be used, in conjunction with the experimental specifications, to reconstruct a path of cell progression. Our algorithm does not make specific assumptions on the path characteristics, e.g. it can be linear or circular. In circular paths any *k*-means cluster can serve as starting point, so that **CONFESS** can use its *M*_*k*_ to reconstruct the path as follows:

1. Find the 2-dimensional average of the centroids, 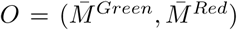, and normalize against it all *M*_*k*_’s so that they are scattered around the center (0, 0) of the Cartesian axis system, 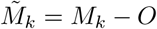 Then, divide the system into its four quadrants;
2. Within each quadrant estimate the tan(*ω*), i.e. the tangent function of the angle between x-axis and the line O*M*_*k*_:

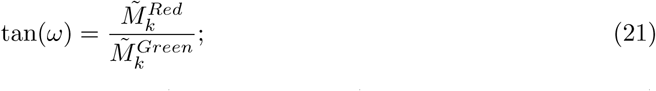
3. Rank the estimates in ascending (clockwise progression) or descending (anti-clockwise progression) order. The ranks define the progression path.

The above methodology works in more general cases, i.e. in paths having single start/end points, where one needs to specify an appropriate starting cluster by viewing the 2-dimensional plot of the log 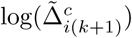 signals.

#### 2.4.2 The Data Driven Haar Fisz transformation for multivariate data

The **CONFESS** model for cell ordering and grouping assumes that the underlying data distribution is approximately normal with constant variance (homogeneity). To meet this assumption, we introduce the novel Data Driven Haar Fisz transformation for multivariate data (*DDHFmv*). Originally, Haar-Fisz (*HF*) has been developed by [35] as a new class of multi-scale, variance stabilization methods to transform 1-dimensional *Poisson*(*µ*)-distributed random variables into *Normal*(*µ*, 1). Data-Driven Haar-Fisz (*DDHF*) extends the above result to a wide range of initial data distributions by modelling the mean-variance relationship as part of the stabilization process [36]. Based on *DDHF*, Motakis et al [37] developed the Data Driven Haar Fisz for single-channel microarrays (**DDHFm**) that outperforms all existing log-based approaches. Here, we show the extension of DDHFm for multivariate data.

##### The mathematical properties of the *DDHFmv*-transform

The *DDHFmv* input vector is the set of the un-logged adjusted fluorescence signals 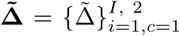 for all cells *I* and channels *c*. Its length is 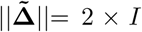. The *DDHFmv* output is a new 2 × *I* vector of approximately normally distributed random variables. The properties of the transform are the following:

1. Property 1: We assume a sequence of independent signals 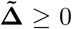 with positive means and variances, 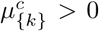 and 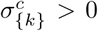, respectively. We require that 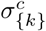 is a non-decreasing function of 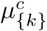, i.e. 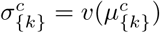 [36]. If *v*() is not known it will be estimated from the data (*data-driven*);
2. Property 2: The 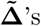 should be sorted by their true cluster means, 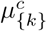, and thus form a piecewise constant sequence. For unknown 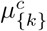 the sorting is done using the observed (estimated) 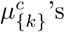. This ordering approximates the true mean ordering in moderate and large samples [37];
3. Property 3: 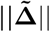 must be a power of two. If not, we reflect the end of the vector in a mirror-like fashion to create artificial data 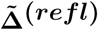 of length -------------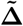----------2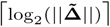 and concatenate them into the new 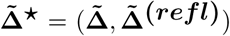 that is a power of two. The *DDHFmv* is applied to the 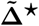 set after which the artificial values can be simply discarded [37].

##### The functions of the *DDHFmv* transform

We present the *DDHFmv* transform of the 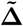 data for *K* clusters and *c* ∈ {*Green, Red*} channels:

1. Denote as 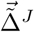 the vector of sorted 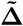 signals (*Properties 2-3*). For example, if *K* = 6 (and ||c||= 2) then 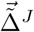 is a concatenated vector of 6 × 2 = 12 sub-vectors, each of variable size *I*_*{k}*_. Examples of sorted sub-vectors can be:

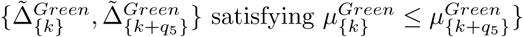

or

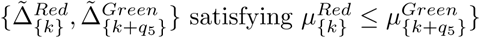

or

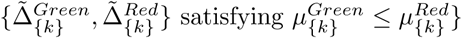

or any other channel / cluster combination for any *q*_5_ so that *k* + *q*_5_ ≤ *K*;
2. For each *j* = *J* - 1, *J* - 2, …, 0 form the smooth and detail coefficients *ξ*^*j*^ and *δ*^*j*^:

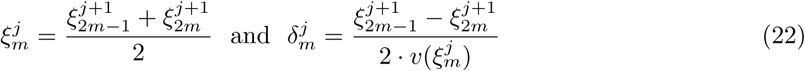

for *m* = 1, …, 2^*j*^ [36];
3. For each *j* = 0, 1, …, *J* − 1 modify the smooth coefficients as:

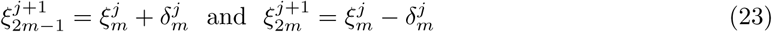

for *m* = 1, …, 2^*j*^. The *ξ*’s of *m* = 2^*j*^ are the *DDHFmv*-transformed data [36];

Following [37], *DDHFmv* estimates the unknown mean-variance relationship, 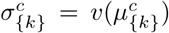 original data via least-squares isotone regression using pool-adjacent violators [38].

### 2.4.3 Iterative scalar projection and change-point analysis

Having collected all computational tools, we are now ready to summarize the iterative approach that performs fluorescence-based cell ordering and phase prediction. The steps are the following:

1. Initial clustering and cell ordering by *k*-means: At the first step we process the 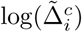 data by (14) - (20) to get the pseudotimes *T*_*i*_ and clusters *K*. These clusters will provide the ordering for the *DDHFmv*-transform;
2. Change-point analysis on the *DDHFmv*-transformed data: To exploit the variance stabilization and gaussianization performance of *DDHFmv*, we use the *k* = 1, *…, K* clusters of the previous step to transform the (un-logged) 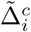 data into *DDHFmv* 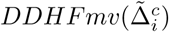. Then for each cell *i* we form the *DDHFmv* counterpart of the log-ratio as:

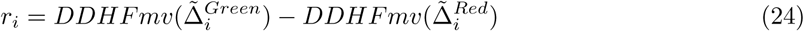

that we sort by *T*_*i*_ in ascending order;
3. The sorted *r*_*i*_’s serve as input for a statistical change point model (e.g. see **ecp**) to estimate significant jumps in the series at a given significance level *α*. The regions separated by the jumps are the updated *k* = 1, …, *K* clusters that reflect significant differences in the relative amounts of 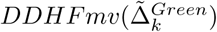 vs 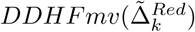 signals across pseudotimes.

Unlike previous flow cytometry oriented models, the above approach enables us to study the spatio-temporal dynamics of a biological process in a versatile, biologically-driven way. In contrast to transcriptome-based statistical models, it functions on the fluorescence data and its results can be integrated with those of single-cell RNA-seq.

## 3 Real and simulated data analysis by CONFESS

In this section we illustrate the use of **CONFESS** on real and simulated datasets coming from Fluidigm single-cell images of the 96 chambers chip. We present its functions and its parameter settings. This paragraph does not cover the important task of biological data interpretation.

### 3.1 The HeLa dataset

The raw data are available for download from ‘SCPortalen: Single-cell centric database’ of RIKEN Center for Integrative Medical Sciences (IMS) [39]. They include 378 images for each of the following sets: Bright Field, Red Channel and Green Channel. The images have been obtained in 5 runs (in this chronological order: 1772-062-248, 1772-062-249, 1772-064-103, 1772-067-038 and 1772-067-039). Each run has been processed at different days except for the last two that they were done in parallel. The scope of the project is to identify the cells on the chip, quantify their fluorescence signal and associate each with a cell cycle phase. Eventually, we wish to study the transcriptional differences across cells of different phases and predict known and novel cell cycle regulators.

#### 3.1.1 Estimation of the spots location

We start by loading **CONFESS**, its dependencies and an indicative subset of the full dataset, **CONFESSdata**. `readFiles()` reads the txt-converted image data. Currently, our technology can directly convert BMP, JPEG and PNG images by specifying their folder at iDirectory. The .C01 conversion tool (https://gitlab.com/al22/r-cellomics) is currently considered for addition in Bioconductor by the respective developers (personal communication with Dr Andreas Leha) and will be added in **CONFESS** in the future. Parameters `BFdirectory` and `CHdirectory` ask for the folder that will contain the BF, Red and Green channel .txt files respectively. If the folder does not exist in the system, it will be automatically generated. The output of `readFiles()` is a list with all file names of interest.

~~~
library("CONFESS")
library("CONFESSdata")
Cpath <-system.file("extdata", package = "CONFESSdata")
files <-readFiles(iDirectory = NULL,
       BFdirectory = paste(Cpath, "/BF", sep = ""),
       CHdirectory = paste(Cpath, "/CH", sep = ""),
       separator = "_", image.type = c("BF", "Green", "Red"),
       bits = 2^16)
~~~

The spot estimation for each image triplet (Section 2.1) is done via `spotEstimator()`.

~~~
result1 <-spotEstimator(files = files,
       foregroundCut = seq(0.60, 0.76, 0.02), BFarea = 7,
       subset = c(), correctionAlgorithm = FALSE)
~~~

The important parameters are `files`, the output of `readFiles()`, that tells the function which file names to analyse; `correctionAlgorithm`, that should be set to FALSE for the first-step spot estimation; and `foregroundCut`, that defines the empirical cutoffs of (1). It is often helpful to train the dataset by picking a subset of well-defined, single-spot images and check the performance using different cutoff values (e.g. the above vs `seq(0.8,0.96,0.02))`. In noisy data we have found that low cutoffs (like the above) produce the best results. Parameter BFarea has two functions: it is both the cutoff of (2) and the size of the rectangular pseudo-object to be measured by BF modelling (Section 2.1). The user can also run the analysis for a subset of the image data, defined by their index number, with the parameter subset. For example:

~~~
result2 <-spotEstimator(files = files,
       foregroundCut = seq(0.60, 0.76, 0.02),
       subset = c(−1, 1:3, 500))
~~~

The above code will ignore indices -1 and 500 as they are non-existent and process only the first three images, and use default values for the other parameters. The association between samples and indices can be inferred from the output variable files above. Intuitively, `subset` can be used to specify the images of a single run but its real usefulness will be described below.

Figure 8 shows the graphical output of `spotEstimator()` for sample 1772-062-248 A12 with an identified spot in both channels (the contour plots on the right panels). The bottom/left panel shows the density of the the channel-specific, background corrected pixel signals. The top/left panel zooms-in the identified location on the BF image. If its central feature is the capture site (as it is the case here), the spot is almost certainly a true cell whose cell cycle phase can be roughly assessed by the densities. Alternatively, the densities can give an insight on the viability of the cell (in visible dead cells they are centered around zero).

**Figure 8:**
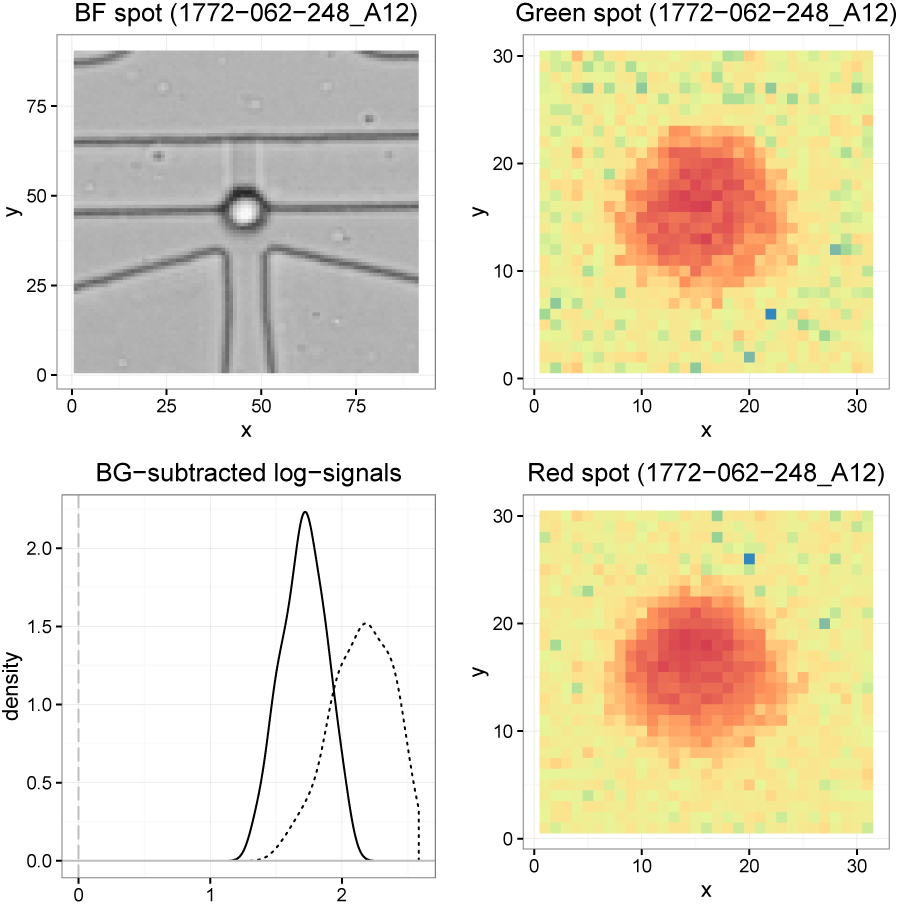
Visualization of the estimated cell of sample ID 1772-062-248 A12. Top left: The estimated cell on the BF image. Right: Contour plot of the estimated cell in each channel. Left/top: The cell location on the original Bright Field image (zoom-in). Bottom left: Background subtracted log-signals.

#### 3.1.2 Quality control and cell signal estimation

**CONFESS** uses visual and statistical inspection tools for the identification of unusually located estimated spots (outliers). The estimates of `result1` are entered into `defineLocClusters()` through the parameter `LocData`:

~~~
clu <-defineLocClusters(LocData = result1, out.method = "interactive.clustering")
Enter the radius of the area containing the reliable coordinates only (suggested values are on the figure):
~~~

The way of processing these data is controlled by out.method whose possible values are: `interactive.clustering` that applies the model of Section 2.1 to flag potential outliers by concentric circles or `interactive.manual` that enables the user to select suspicious data manually by point-and-click on the plot (see R function `locator()`). Figure 5 showed previously the characteristic concentric circles indicating outliers at different significance levels (the cyan circle is constructed at *α* = 5%). The user is required to enter the radius of a circle which will flag all dots that lie on or outside it. The main component of the output variable clu is the previous spot location matrix augmented with an extra outlier index.

The re-estimation step requires a second round implementation of `spotEstimator()` with `correctionAlgorithm = TRUE` and the attention shifted to a different set of parameters: `subset` inputs the outlier sample indices, QCdata reads the spot location estimates, including the capture site directionality and the 2-dimensional medians by run and chamber ID. Where BF modelling is not accurate, the algorithm sets as spot center the median location of all estimated images of a particular run / chamber direction (`median.correction`).

~~~
resultOut <-spotEstimator(files = files, subset = clu$Outlier.indices,
       foregroundCut = seq(0.60, 0.76, 0.02),
       correctionAlgorithm = TRUE,
       QCdata = clu, median.correction = TRUE)
~~~

Before producing the final cell location and raw signals matrix, the user can re-assess the accuracy of the second round estimates by `defineLocClusters()`. Then, the output of interest is obtained via `LocationMatrix()` that, apart from listing the locations and the signal estimates, quantifies the cell’s existence and viability. This is done by setting appropriate filters at filter.by. Here, we only keep the ‘confidence’ estimates (non-rejected locations in Figure 5) that reject 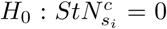 at FDR = 0.5%. textbfCONFESS offers a wide selection of other filters such as the spot area, the fluorescence signal and the number of secondary spots.

~~~
clu <-defineLocClusters(LocData = resultOut,out.method = "interactive.manual")
Results <-LocationMatrix(data = clu,
       filter.by = matrix(c("FDR", "Out.Index", 0.005, "confidence"), ncol = 2))
~~~

Table 1 shows the tabulated output of the first three samples. The first column contains the sample ID and the next three summarize the identified location (3) - (4). The raw fluorescence characteristics of each channel are described by the average foreground and background signals in the middle part of Table 1 that also contains the signal-to-noise ratio estimates (7) and the assessment of cell existence by statistical testing. The bottom part of Table 1 shows the confidence index (non-rejected locations in Figure 5; alternative is ‘outlier’), the number and the location of secondary spots and the existence or not of a true cell. For sample 1772-062-248 A03 we found one extra spot in the Green channel (X = 262, Y = 368) that was considered an outlier. The spot at X = 262, Y = 368 in the Red channel had a strong signal on the appropriate location.

**Table 1:** Cell location estimates by **CONFESS**.

| SampleID | X | Y | Size | Estimation.Type |  |  |
| --- | --- | --- | --- | --- | --- | --- |
| 1772-062-248_A01 | 259 | 367 | 31 | Fluorescence-based |  |  |
| 1772-062-248_A02 | 261 | 335 | 49 | Chip.Pattern-based |  |  |
| 1772-062-248_A03 | 262 | 368 | 19 | Fluorescence-based |  |  |
| fore.Green | back.Green | fore.Red | back.Red | Signal-to-Noise | Pvalue | FDR |
| 48.399 | 17.173 | 219.071 | 17.290 | 3.561 | 6.1e-7 | 1.7e-6 |
| 18.352 | 16.756 | 19.771 | 17.863 | 0.142 | 9.9e-1 | 1.0e+0 |
| 26.184 | 16.966 | 86.503 | 17.005 | 2.224 | 7.1e-5 | 1.8e-4 |
| Out.Index | Other.Spots |  |  | Cells |  |  |
| Confidence | 0 |  |  | 1 |  |  |
| Confidence | 0 |  |  | 0 |  |  |
| Confidence | X = 30, Y = 204 (Green) — X = 262, Y = 368 (Red) |  |  | 1 |  |  |

The above metrics and image outputs are indicators of the data quality. **CONFESS** offers valuable information for the difficult and time-consuming task of single-cell quality control that we suggest to always combine with thorough single-cell RNA-seq data analysis [4] in order to retrieve a reliable list of cell estimates. A first step to this end is to collect **CONFESS**’s reliable estimates with:

~~~
Reliable.results <-Results[which(as.numeric(Results$Output$Cells) == 1), ]
~~~

The automated image analysis estimates of Table 1 were coupled with manual check of the output plots to identify 287 out of 378 samples with a single detected cell on the capture site. We report that 317 out of 378 images, independently of whether they carry one cell, multiple cells or nothing, are of acceptable quality (visually in-focus). Extensive analysis using the above pipeline and scRNA-seq data modelling led to the identification of 246 reliable cells. The integration of image and scRNA-seq data is beyond the scope of this paper and it is not presented. Indicatively though, 83.8% of **CONFESS**’s predictions were confirmed after the integration.

#### 3.1.3 Fluorescence signal adjustment

To perform signal adjustment we collect the data of the 246 reliable samples of the previous step as:

~~~
step1 <-createFluo(from.file = system.file("extdata","Results_of_image_analysis.txt", package = "CONFESS"), separator = "_")
[1] "Run 1772-062-248 was converted into 1"
[1] "Run 1772-062-249 was converted into 2"
[1] "Run 1772-064-103 was converted into 3"
[1] "Run 1772-067-038 was converted into 4"
[1] "Run 1772-067-039 was converted into 5"
~~~

data contains the estimates in the format of Table 1. Alternatively, one could point to a data file (from.file) that would force the algorithm to ignore the input of data parameter. In this way, the adjustment and all subsequent analytic steps can be independent of our image analysis results. In any case, step1 will store the sample IDs, the raw fluorescence signals, the cell sizes and the Run IDs of all reliable samples into a list. We can quickly check the distribution of the *normexp* adjusted signals (9) by run (parameter batch) using the following commands:

~~~
step2a <-FluoSelection_byRun(data = step1, batch = 5)
step2a <-getFluo_byRun(data = step2a, transform = "log")
~~~

The first line selects the data of run ID 1772-067-039 (batch = 5) which `getFluo byRun()` subsequently uses to generate the log-transformed background corrected signal densities (Figure 6). The densities highlight the differences across channels and help us select the candidate run adjustment models (11) - (13). The data indicate the existence of at least two modes in each channel. Similar plots are obtained for the other runs, so that the adjustment will be based on finite mixtures of regression models.

~~~
step2b <-Fluo_adjustment(data = step1, transform = "log", maxMix = 3,
       prior.pi = 0.1, single.batch.analysis = 5, seed = 999)
step2b <-getFluo(data = step2b)
~~~

The model selection is done by `Fluo_adjustment()` that fits the log-transformed raw signals in (10) - (12) with *M*^*c*^ = 3 `(maxMix)` and *π*_*m*_^*c*^ = 0.1 `(prior.pi)`. The data are run adjusted using the fifth run (1772-067-039) as baseline `(single.batch.analysis)` and will be subsequently *normexp* corrected. Figure 9 shows the uncorrected signals (top) that have been subsequently corrected (bottom) by *normexp* and **flexmix** with 2 components. For reproducibility reasons the user has the option to specify a seed number (seed) for the **flexmix** MC sampling (default is NULL). Next, `getFluo()` stores the corrected signals into step2b that serves as input for unsupervised cell ordering and clustering. For data coming from a single run, both `Fluo_adjustment()` and `getFluo()` are replaced by `getFluo_byRun()` that calls the *normexp* model directly.

**Figure 9:**
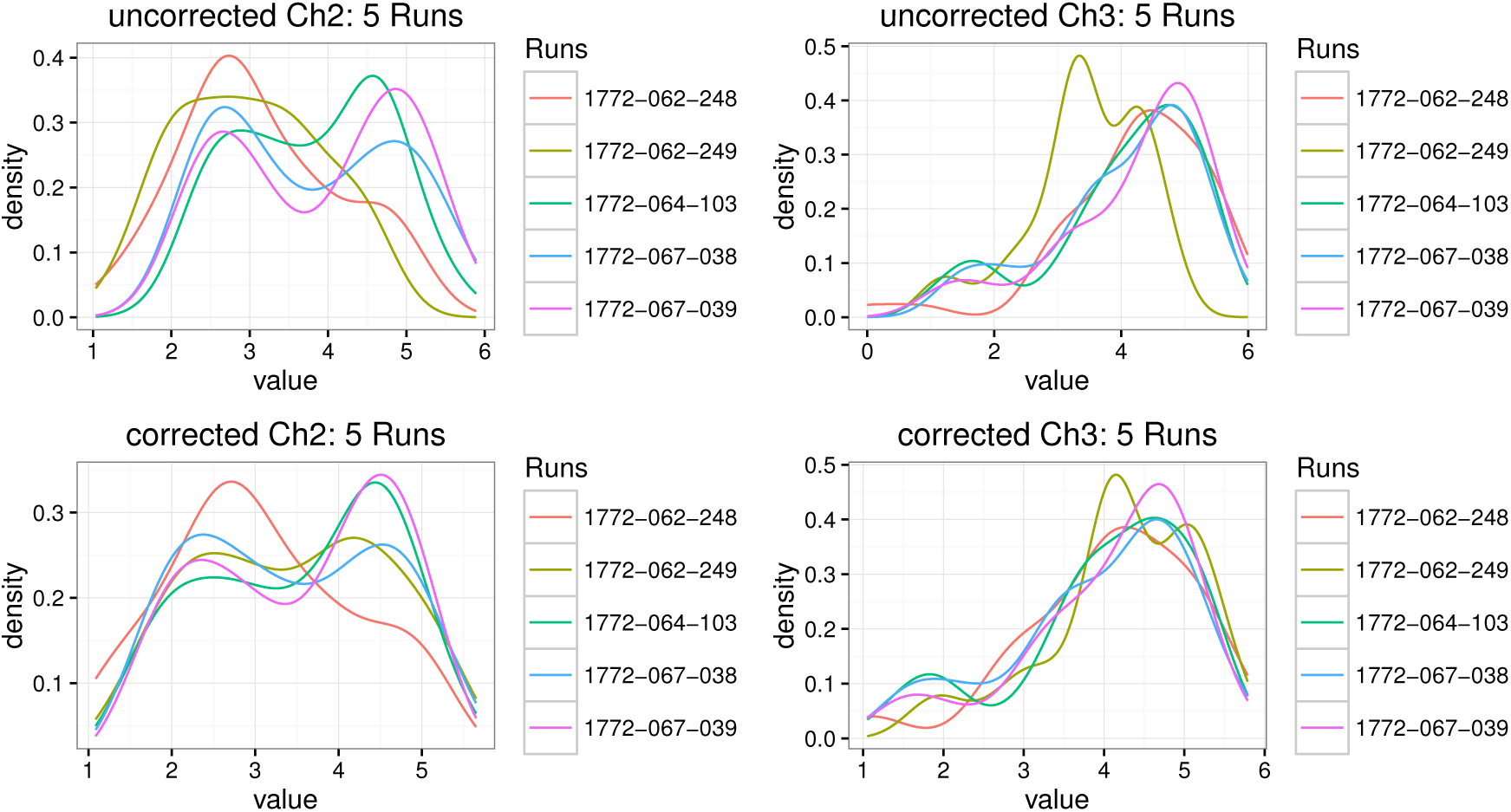
Background adjustment by normexp and run effect correction by **flexmix** of the channel signals of each run. Top: uncorrected channel signals. Bottom: corrected channel signals. The colors indicate the different runs.

#### 3.1.4 Unsupervised cell ordering and grouping

The adjusted data are loaded for path progression estimation (14) - (21) as:

~~~
step3 <-Fluo_inspection(data = step2b, altFUN = "kmeans", k.max = 15)
step3 <-pathEstimator(step3, path.start = 3,path.type = c("circular", "clockwise"))
~~~

The initial clustering and their centroids (14) are controlled by altFUN (*k*-means is the default method) and k.max specifying the maximum number of clusters. The clusters estimate the progression path (21) in `pathEstimator()` for any arbitrary starting cluster `(path.start)` and directionality type (path.type). A circular path type implies circular-like progression in contrast to A2Z that is used for paths with well-defined start and end points (e.g. in cell differentiation) or other for arbitrary path types with unclear directionality (clockwise or anti-clockwise). If path.type = ‘other’, the user is asked to manually define the progression path by examining an informative figure (Figure 7 obtained by `Fluo_inspection()`). The cyclic progression path can be automatically estimated in `pathEstimator()` as 3-4-6-2-1-5.

~~~
step4 <-Fluo_modeling(data = step3, init.path = step3$Path,
       VSmethod = "DDHFmv", CPmethod = "ECP",
       CPpvalue = 0.01, CPmingroup = 10)
step5 <-Fluo_ordering(data = step4, den.method = "wavelets")
~~~

The iterative unsupervised cell ordering and grouping is performed by `Fluo_modeling()` using either *DDHFmv* or log transform (`VSmethod`) and *ECP* or *PELT* (CPmethod) for change-point analysis at a given significance level (`CPpvalue`). Here, the user-defined parameter CPmingroup specifies that a change-point estimated cluster is accepted if it contains at least 10 cells. If the user believes that the above estimated path is incorrect, he could manually replace it by his own choice in `init.path`. Next, `Fluo_ordering()` generates the output tables and plots. It is only coincidence that the number of groups in Figure 7 and Figure 10 are the same. The cluster structures are substantially different.

**Figure 10:**
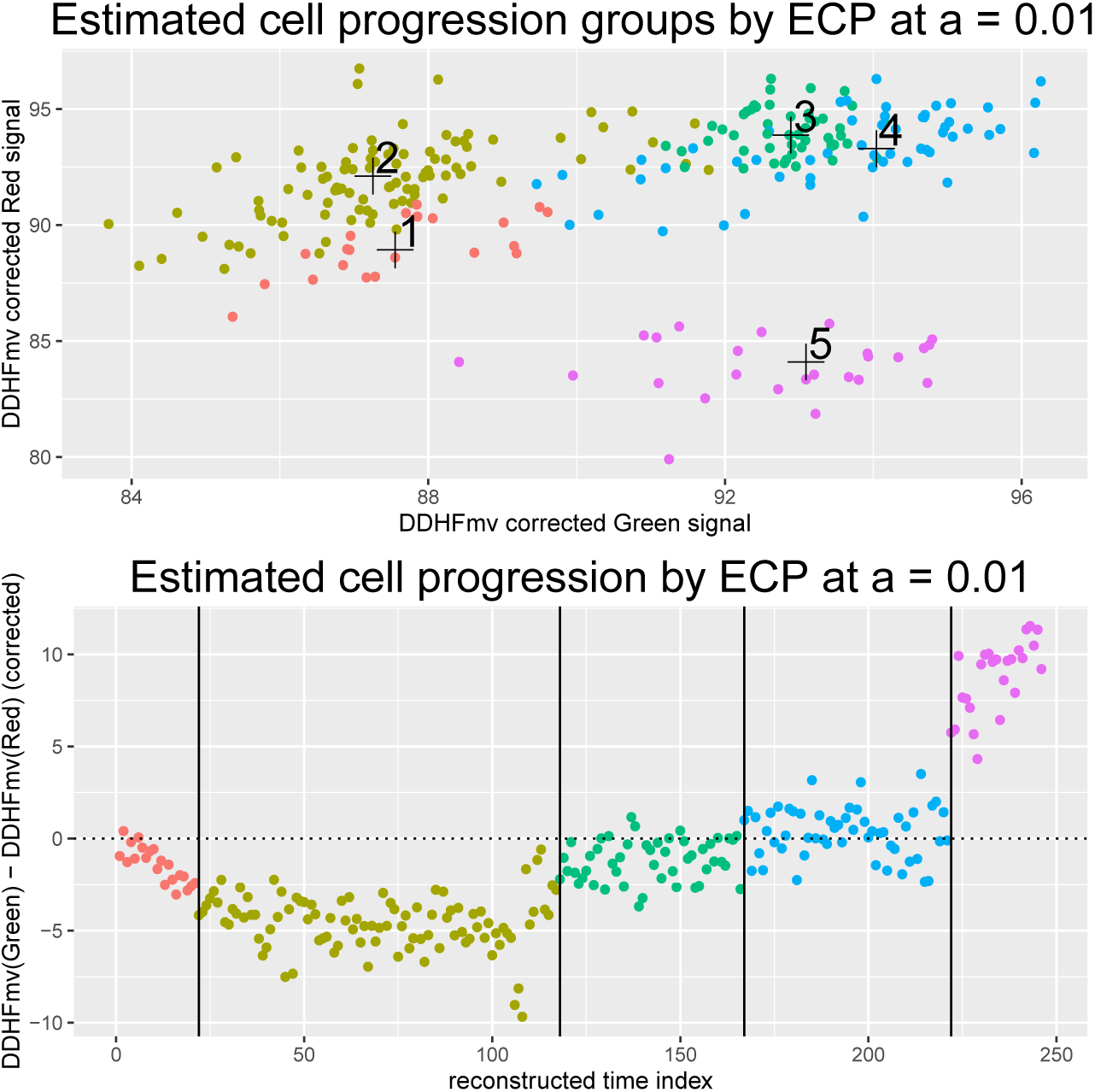
Top: Scatterplot of the *DDHFmv*-transformed corrected signals indicating the estimated clusters (colors). Bottom: Pseudotime series plot of the difference between the *DDHFmv*-transformed corrected signals indicates the predicted cell sorting within and across clusters (the colors match the ones of the top panel). The vertical lines show the estimated change-points by **ecp** at *a* = 1%.

Figure 10 shows the final groups in the 2-dimensional plane (top) and the cell ordering by pseudotime (bottom). The first lines of the output table are depicted in Table 2. The residuals are estimated as

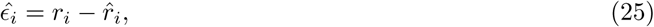

where the *r*_*i*_’s come from (24) and 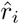 *r* are the wavelets fitted *r*_*i*_’s (parameter `den.method`). The output also includes residual diagnostic plots (histogram) and metrics (normality tests). The Grubbs statistic can be used on the 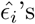 to flag outlying cells at *α* = 1%.

**Table 2:** Fluorescence signal estimates, pseudotimes and clusters (cell cycle phases) by **CONFESS**.

| Index | SampleID | Runs | Size | Corr.Size | Pseudotime |
| --- | --- | --- | --- | --- | --- |
| 1 | 1772-062-248_A01 | 1772-062-248 | 31 | 3.43 | 110 |
| 2 | 1772-062-248_A03 | 1772-062-248 | 19 | 2.94 | 58 |
| 3 | 1772-062-248_A04 | 1772-062-248 | 152 | 5.02 | 229 |
| $\log(Red)$ | $\log(Green)$ | $DDHFmv(Red)$ | $DDHFmv(Green)$ | | |
| 3.28 | 5.25 | 90.19 | 94.86 |  |  |
| 2.25 | 4.23 | 86.28 | 92.48 |  |  |
| 3.21 | 1.84 | 88.41 | 84.10 |  |  |
| $DDHFmv(Red) - DDHFmv(Green)$ | | | Clusters | Residuals | Out.Index |
| -4.67 |  |  | 2 | -0.79 | normal |
| -6.19 |  |  | 2 | -1.15 | normal |
| 4.31 |  |  | 5 | -1.85 | normal |

Figure 10 shows a clear cell cycle progression path. Following the logic of the Fucci system, we can infer that the first small cluster 1 highlights cells at the end of mitosis (EM) and early G1. The vast majority of HeLa cells are found in the next phases, i.e. G1, that is represented by cluster 2, late G1 - to early S (cluster 3) and S (cluster 4). The remaining cells have been estimated in G2.

In agreement with previous findings on HeLa, **CONFESS** predicted that 83% of the cells are found in G1 and S phases whose duration is expected to be substantially long [40]. The model residuals (25) were found to be approximately normally distributed (Agostino test P-value for skewness = 0.093; Bonett test P-value for kurtosis = 0.874; KS-test P-value for Normality = 0.585; Jarque test P-value for Normality = 0.247) implying that our wavelets non-parametric curve fitting model can appropriately detect the significant jumps across pseudotimes.

### 3.2 Data simulation

To test the accuracy of our cell location estimates we have built a data simulator function for Fluidigm C1 two-channel experiments. Its various parameters are described below:

~~~
simresult <-simcells(channels = 2, spots.per.image = c(2, 1),
       image.dimensions = rep(100, 2), agreement.number = 1,
       signal.level = list(c(700, 400), 700), noise.level = rep(100, 2),
       spot.size = list(c(30, 20), 30), one.location = c(50, 50))
~~~

The presented demonstration of simcells() generates Case 2A of Figure 2. The two images (channels) correspond to pseudo-Green containing two spots (spots.per.image) and pseudo-Red with one spot, respectively. The image dimensions are 100 × 100

~~~
(image.dimensions).
~~~

Only one matching pair is simulated (`agreement.number`) whose location on both images is defined by one.location (the center of the image). The spot sizes (`spot.size`) and signals (`signal.level`) enter in lists whose length (components) equal to the number of spots in each channel. The first element of each component always corresponds to the matched pair. The channel-specific noise level is defined in noise.level. Thus here, the pixel signals of the matching spots of size 30 pixels are both drawn from a *Poisson*(700)+*Normal*(*µ*^*c*^, 100) where *µ* is internally calculated so that the minimum pixel signal of channel *c* image is non-negative.

Function `simcells()` can generate multiple spots of various sizes, signals and on different locations. More than one matching pair can be generated (e.g. case 3B of Figure 3) by setting agreement.number =

The output includes a graphical representation of the spots on the two channels and a matrix of their locations with their average noisy fluorescence signal. In this framework, we test **CONFESS**’s ability for cell identification. We simulate 317 two-channel images of dimensions 512 × 512 from the following settings:

1. Low-noise data: Each channel contains at least one or multiple spots. There is only one matching pair and its center is found in the location (*X*^*\**^, *Y* ^*\**^) on both channels (simulating a true cell). The rest of the spots are noise and should not be matching. The spot pixel signals for each spot *s*, image *i* and channel *c* are drawn from a (*μ*_1_) + 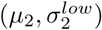. Each of these parameters varies randomly with *µ*_1_ ∈ [200, 1400] and 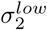 ∈ [100, 200] (*µ*_2_ is internally estimated). Low *µ*_1_ values correspond to weak spots in one or both channels. The spot sizes are drawn from [20, 250]. The spot center (*X*^*\**^, *Y* ^*\**^) is derived by manual identification of the capture site of the 317 acceptable quality BF images of the real dataset. Thus, if fluorescence estimation fails **CONFESS** will interrogate the BF data;
2. High-noise data: We keep the above randomly selected central locations, spot sizes, spot signals and spot numbers but we define 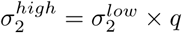 with *q* drawn at random from *q* ∈ [2, 2.5];
3. Negative controls: In this set we do not simulate any spots. The noise of the 317 two-channel images is simulated from *Normal*(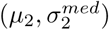) with 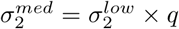 and *q* ∈ [1, 2]. This scenario resembles the case of empty images or images with weak spots in both channels. **CONFESS** should rely on the BF data for estimation of the capture site. We would like to test how successful we are in recovering it;
4. Contaminated data (double spots): Each channel contains exactly two spots with matching locations implying image contamination. One spot pair is always located at (*X*^*\**^, *Y* ^*\**^) on both channels while the other can be anywhere in the image. The spot signals are generated as 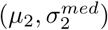. This setting evaluates how well **CONFESS** identifies contaminated data to discard them in the appropriate time.

Below, we provide an small example of the actual code for the first 3 images of the low-noise scenario. The high-noise data can be easily obtained by changing the noise parameter (see below). The negative controls are obtained by setting the number of spots for each channel (spots.per.image) to be equal to zero. The double spots are similarly generated by setting the number of spots for each channel to be equal to two.

First, we store the central location of the capture sites (it has been determined by manual inspection):

~~~
cen <-matrix(cbind(c("1772-062-248_A01", "1772-062-248_A02", "1772-062-248_A03"),
       c(259, 259, 261), c(367, 341, 370)), ncol = 3)
~~~

Then we randomly assign a number of spots (0 to 3) in each channel. Note that in the low-noise and high-noise scenarios we do not allow the images to be ‘empty’. When zero is picked, we force the algorithm to generate one weakly expressed spot.

~~~
sp <-matrix(sample(0:3, (2 * nrow(cen)), replace = TRUE), ncol = 2) w <-which(apply(sp, 1, sum) == 0)
if (length(w) > 0) {
          for (i in 1:length(w)) {
                  sp[w[i], ] <-c(1, 0)
          }
}
~~~

Next, we set the signal, the noise and the size parameters. Note that the signals of the low-noise are well above the noise levels. However, when we draw a ‘zero’ for a channel the signal falls near the noise level.

~~~
signals <-seq(800, 1400, 1)
sizes <-seq(20, 250, 1)
noises <-seq(100, 200, 1)
~~~

We prepare the output variables:

~~~
sp.alt <-sp
resultCh2low <-resultCh3low <-matrix(0, nrow(sp), 6)
~~~

Finally, we generate the spots and signals for each channel:

~~~
for (i in 1:nrow(cen)) {
         noi1 <-sample(noises, 1)
         if (sp[i, 1] > 0) {
                 sig1 <-sample(signals, sp[i, 1])
                 siz1 <-sample(sizes, sp[i, 1])
         } else {
         }
                 sp.alt[i, 1] <-1
                 sig1 <-round(noi1 * sample(seq(2, 2.5, 0.05), 1), 0)
                 siz1 <-sample(sizes, 1)
         }
         noi2 <-sample(noises, 1)
         if (sp[i, 2] > 0) {
                 sig2 <-sample(signals, sp[i, 2])
                 siz2 <-sample(sizes, sp[i, 2])
         } else {
         }
                 sp.alt[i, 2] <-1
                 sig2 <-round(noi2 * sample(seq(2, 2.5, 0.05), 1), 0)
                 siz2 <-sample(sizes, 1)
         }
         siz1[1] <-siz2[1]
         siz1[1] <-round(siz1[1] * sig1[1] / sig2[1], 0)
         r <-simcells(channels = 2, spots.per.image = sp.alt[i, ],
                 one.location = as.numeric(cen[i, 2:3]),
                 image.dimension = rep(512, 2),
                 signal.level = list(sig1, sig2),
                 noise.level = c(noi1, noi2),
                 spot.size = list(siz1, siz2), agreement.number = 1)
         resultCh2low[i, ] <-c(cen[i, ], sp.alt[i, 1],
                 as.numeric(r$Spots$Ch1[1, 3]), siz1[1])
         resultCh3low[i, ] <-c(cen[i, ], sp.alt[i, 2],
                 as.numeric(r$Spots$Ch2[1, 3]), siz2[1])
}
~~~

Note that this exercise could also serve as an example of **CONFESS**’s application on data from a single run where cell ordering is not needed. Such type of analysis could be used to link the cell estimates to a given experimental design, e.g. the cells with 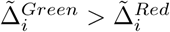 come from a hypothetical *condition1* and the rest from a hypothetical *condition2*.

#### 3.2.1 Discussion of the simulated data results

We evaluate the accuracy of **CONFESS** in predicting the cell location, the cell size and the cell signal in both channels. To this extent, we calculate the following quantities:

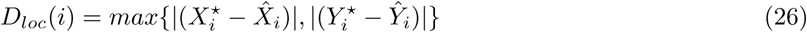

that assesses the location estimation error in pixels for each cell *i*,

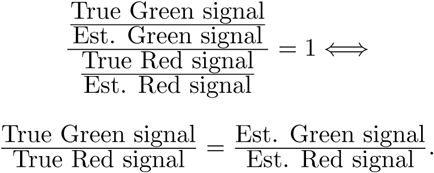

Estimated size that assesses the size estimation error in pixels for each cell *i* and

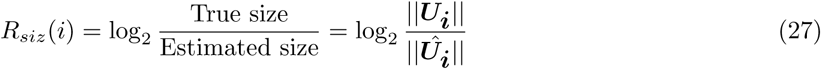

that performs a cross channel signal comparison to identify possible biases in the estimation. We wish to show that the channel specific signal estimation errors are comparable irrespectively of how big the errors are (which strongly depends on the noise level, the signal level and the cell size). Comparable errors imply that none of the channel signal is over / under estimated compared to the other which can seriously affect the data inference. To assess the average error rate of (26) - (28) we run **CONFESS** on 10 simulated datasets (generating in total 3,170 cells) from each of the above categories.

Figure 11 (top) shows the absolute location differences (26) of the low and high noise simulated samples for the first iteration. The vast majority of the samples have very low location errors. The superimposed dots highlight samples for which **CONFESS** failed to identify a cell (*Cells* = 0 in Table 1). There are 4 such samples in the Low-noise (discovery error = 1.2%) and 12 such samples in the High-noise (discovery error = 3.7%).

**Figure 11:**
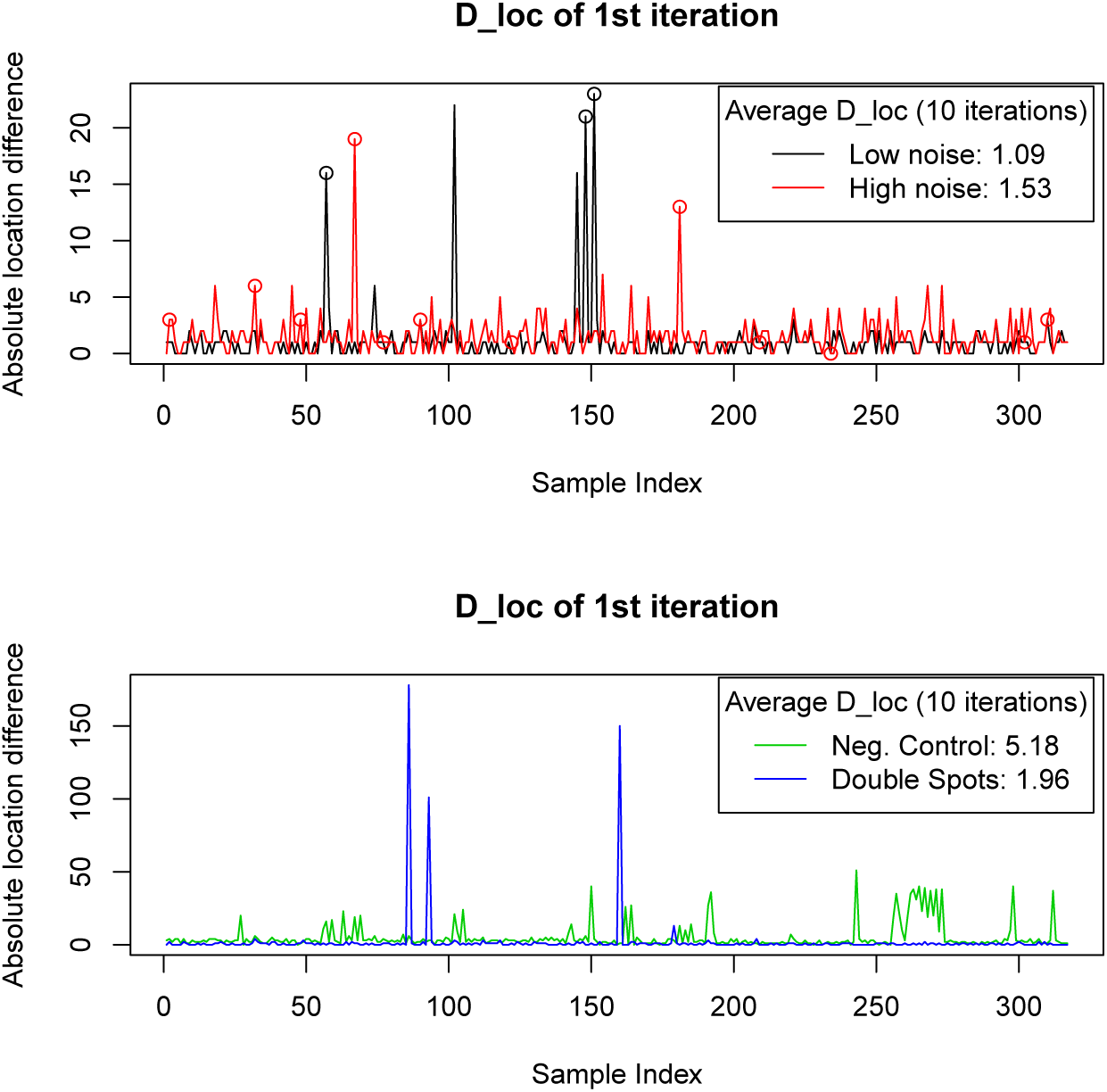
Absolute differences between the estimated and the true cell locations of the simulated data for the 1st iteration. Top: Results of the low and high noise simulated datasets. Bottom: Results of the negative control and contaminated datasets. The dots highlight the cases that have been automatically flagged having zero cells. The legends indicate the average error rate in all 10 iterations.

Among the former there is one case (sample index 90 that is also missed in the High-noise dataset) for which the cell location has been accurately found (*D*_*loc*_(90) = 3) but the signal was very low in both channels to be detected and thus filtered out by our filters (filter.by). Manual image inspection (quality control) back-retrieved the missing information, thus dropping the discovery error to 0.09%. Similarly, among the High-noise missing data, 9 of them exhibit *D*_*loc*_(*i*) ≤ 3 and can be found in the images but their signal was very low in both channels to be detected. As before, the actual discovery error is as small as 0.09%.

Figure 11 (bottom) shows the *D*_*loc*_’s of negative controls and contaminated datasets that exhibit substantially higher error rates. In the case of negative controls our estimates assess how successfully **CONFESS** located the capture site. In approximately 90% of the cases the site was predicted to be less than 5 pixels away from the truth which implies that we estimate accurately the capture site (it architecture resembles a circular-like region of radius 5 pixels). In all cases **CONFESS** has accurately predicted the cell absence, i.e. it does not report artificial cell-like regions.

The contaminated data simulation is informative for two reasons: first, we test **CONFESS**’s ability to identify contaminated images that in any real data study should be filtered out; second, we examine the accuracy of the location prediction. Our methodology identified the contaminated data in 99.3% of the cases (315 simulated samples out of 317).

Table 3 shows the average error rates of 10 simulated data sets of 317 samples. In both low and high noise data, the average absolute displacement of the cell’s center is very low, averaging to 1.09 and 1.53 pixels respectively. Considering that each simulated cell is at least 20 pixels large, this result implies that we perform a very successful estimation in both cases. Similar low errors are obtained for the contaminated data (1.96 pixels) indicating the **CONFESS** can reliably find the true cell. Naturally, the negative controls show a higher error rate since the algorithm fully relies on BF image modelling to locate the capture site which is on average 5.18 pixels away from the manually retrieved ‘ideal’ location.

**Table 3:** Average error rates of 10 simulated datasets of 317 samples each.

|  | Low-noise | High-noise | Neg. controls | Contaminated |
| --- | --- | --- | --- | --- |
| Absolute $D_{loc}$ | 1.09 | 1.53 | 5.18 | 1.96 |
| Original (absolute) $R_{siz}$ | -0.07 (0.20) | 0.10 (0.22) | — | — |
| Original (absolute) $D_{sig}$ | 0.06 (0.16) | 0.07 (0.19) | — | — |

We calculated the average (and absolute) *R*_*siz*_ which quantifies the log difference between the true and the estimated cell size (obtained by (4)). The values outside the parentheses in Table 3 show that in the Low-noise dataset the ratio of true vs estimated sizes (in pixels) is 2^-0.07^ = 0.95 and thus we tend to slightly underestimate the true spot sizes. The opposite is true for the High-noise data with average ratio 2^0.1^ = 1.07. We evaluated the differences irrespectively of direction (absolute differences) to find ratios of 2^0.20^ = 1.14 and 2^0.22^ = 1.16 for each dataset, respectively. The discrepancy of the Low- and High-noise estimates occur because we used the same textttforegroundCut on both datasets. The values imply that we miss the size of the true object by approximately 15%. Put simply, for an object of size 100 pixels our estimation is averaging to 85 or 115 pixels. For rectangular shapes (10 pixels per side) this would roughly mean that we remove or add one layer of pixels from one side.

Finally, we compare the ratios of true vs estimated signal across channels by *D*_*sig*_. A log ratio of 0 (or a ratio of 1) indicates that **CONFESS** can consistently approximate the true relationship between the signals of the two channels since:

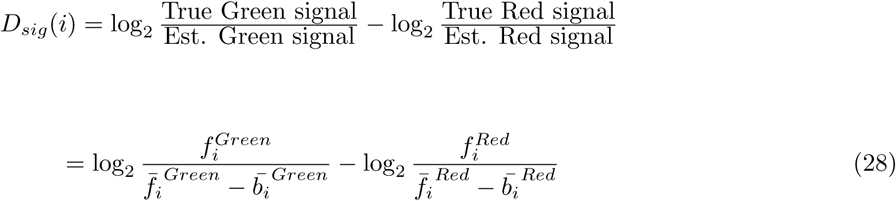

Evidently, both the original (2^0.06^ = 1.04 and 2^0.07^ = 1.05) and the absolute differences (2^0.16^ = 1.11 and 2^0.19^ = 1.14) are relatively low in the two datasets. Summarizing the above results we conclude that **CONFESS** performs reliable cell identification analysis for a wide range of signal, noise and size parameters in different scenarios (positive / negative controls and contaminated data).

## 4 Robustness of CONFESS predictions

To computationally evaluate CONFESS predictions we apply the following cross - validation scheme:

1. Remove all samples of Run 1 and generate new *k*-mean clusters;
2. Put back the excluded samples and re-estimate them (based on the new clusters);
3. Estimate the cross-validated pseudotimes and phases. For each sample *i*, compare the original pseduotimes *T*_*i*_ against the cross-validated pseudotimes 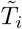 as 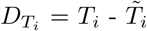
4. Repeat the above steps for all runs b = 2, …, 5. Thus, in each iteration we remove / re-estimate a different number of samples (among the reliable of step1 of Section 3.1).

**CONFESS** uses the functions `Fluo_CV_prep()` and `Fluo CV modeling()` to apply the above pipeline and assess the error of the location estimates by sample:

~~~
step1 <-createFluo(data = Reliable.results, from.file = FALSE)
steps2_4 <-Fluo_CV_prep(data = step1, init.path = rep("bottom/left", 2),
       path.type = c("circular", "clockwise"), maxMix = 3,
       single.batch.analysis = 5, prior.pi = 0.1,
       VSmethod = "DDHFmv", CPmethod = "ECP", CPpvalue = 0.01)

co <-combn(5, 4)
steps2_4cv <-Fluo_CV_modeling(data = steps2_4, B = 1, batch = co[, 1],
       seed.it = TRUE)$All.Progressions[[1]]

for (i in 2:ncol(co)) {
       steps2_4cv <-matrix(cbind(steps2_4cv,
              Fluo_CV_modeling(data = steps2_4, B = 1, batch = co[, i],
              seed.it = TRUE)$All.Progressions[[1]][, 2]),
              nrow = nrow(steps2_4cv))
}
dd <-abs(apply(steps2_4cv, 1, function(x) mean(x[1] - x[2:length(x)])))

barplot(dd, col = factor(steps2_4$Batch5$batch),
       xlab = "Samples (sorted by run)",
       ylab = "Average difference: Original pseudotime - CV(b) Pseudotime",
       sub = "The dashed line at 1 is the median error")
       abline(h = ceiling(median(dd)), lty = 2)
~~~

Figure 12 shows the average difference of each sample over all iterations. The values are generally low (the median error is 1) apart from those of 15 samples that exhibit relatively high variability in their predicted pseudotimes. The analysis enables us to discover such outliers and study them further.

**Figure 12:**
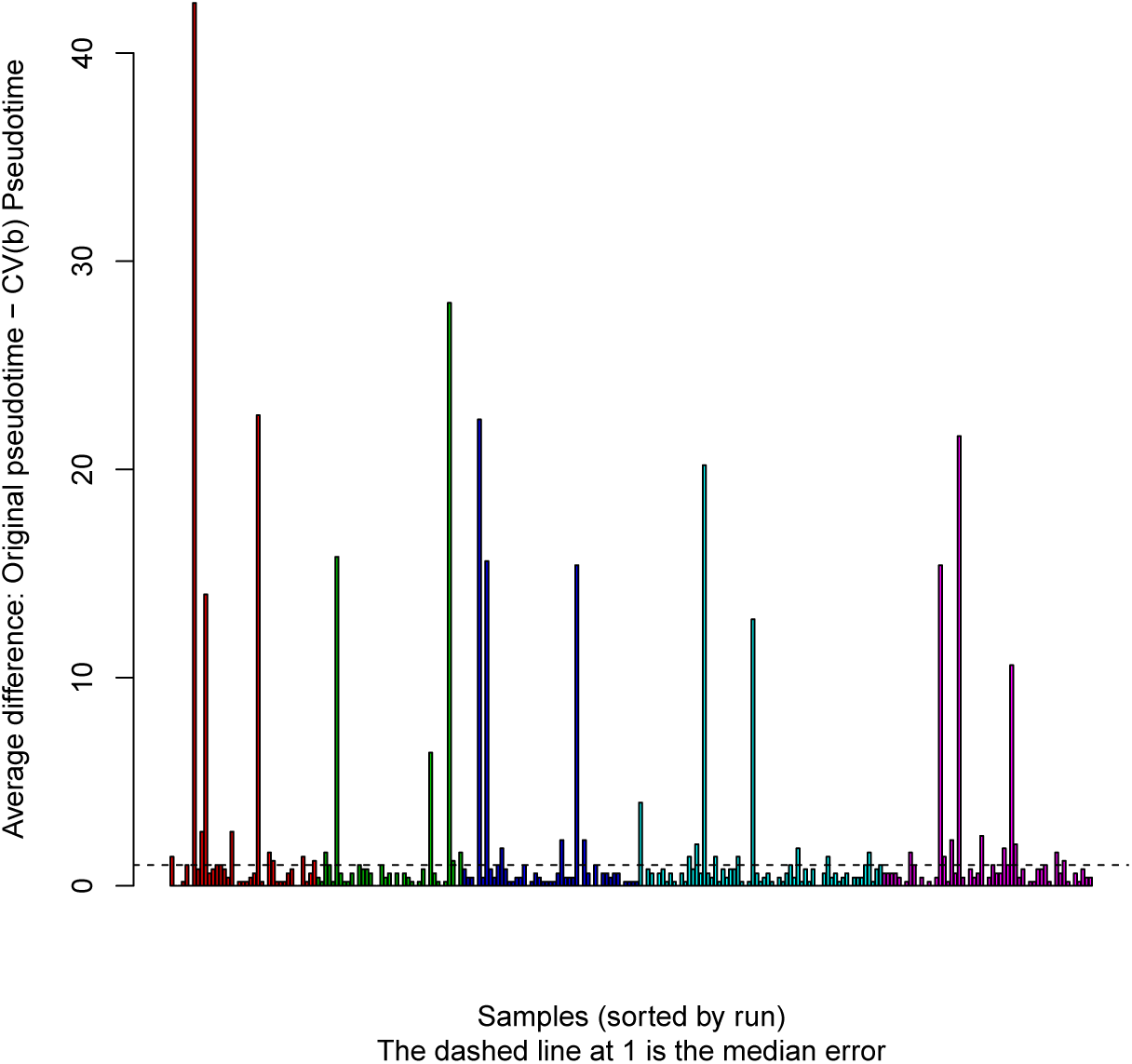
Evaluation of location errors by **CONFESS**’s 5-fold cross-validation.

## 5 Comparison of CONFESS to existing approaches

The unsupervised clustering of fluorescence signals has been previously discussed in the flow cytometry literature. Peel and McLachlan [41] utilize expectation-conditional maximization to fit multivariate *t*-mixture models and predict a set of clusters without relying on spherical shapes (**flowClust**). **SamSPECTRAL** [21] utilizes spectral clustering and graph theory to reach an optimal solution. **flowMerge** [22] estimates the optimal number of those clusters by the Bayesian Information Criterion. **flowMeans** [23] and **flowPeaks** [24] use *k*-means to produce a set of starting clusters that are iteratively refined by change-point analysis and smoothing respectively. We re-estimate the cell cycle phases (clusters) by each of the above models (**flowMerge** is an extension of **flowClust**) and compare their results against **CONFESS**. The flow cytometry results can be directly obtained from **CONFESS** as:

~~~
mod.fm <-Fluo_inspection(data = step2b, altFUN = "fmerge", savePlot = getwd())
length(which(mod.fm$GAPgroups[, 2] == 2))
[1] 9

mod.peaks <-Fluo_inspection(data = step2b, altFUN = "fpeaks", savePlot = getwd())
length(which(mod.peaks$GAPgroups[, 2] == 2))
[1] 27

mod.peaks.nooutliers <-FluoSelection_byRun(data = mod.peaks,
       other = which(mod.peaks$GAPgroups[, 2] != 2))

mod.sam <-Fluo_inspection(data = step2b, altFUN = "samSpec", savePlot = getwd())

mod.means <-Fluo_inspection(data = step2b, altFUN = "fmeans", savePlot = getwd())
~~~

Figure 13a-13b show that **flowMerge** and **flowPeaks** generate 3 main clusters that separate G1, S and G2 -like phases. Highlighted by ‘-999’ are two small clusters of 9 and 27 samples respectively that the algorithms themselves flag as outliers. **SamSPECTRAL** gives a similar solution with large G1 and S clusters and a smaller G2 (Figure 13c). Cluster 4 seems to be an intermediate between S and G2 consisting of 9 samples. Cluster 5 could be considered as an outlier set of size 3 but it does not marked as such. The worst predictions are obtained by **flowMeans** that returns a single cluster (Figure 13d). By design, none of the above methods can reconstruct the spatio-temporal progression of the cell cycle or of any other biological system, nor it provides enough data towards this goal. **CONFESS** can essentially be used in conjunction with any of these clustering algorithms to predict a given progression path and facilitate the deeper understanding of a given biological mechanism.

**Figure 13:**
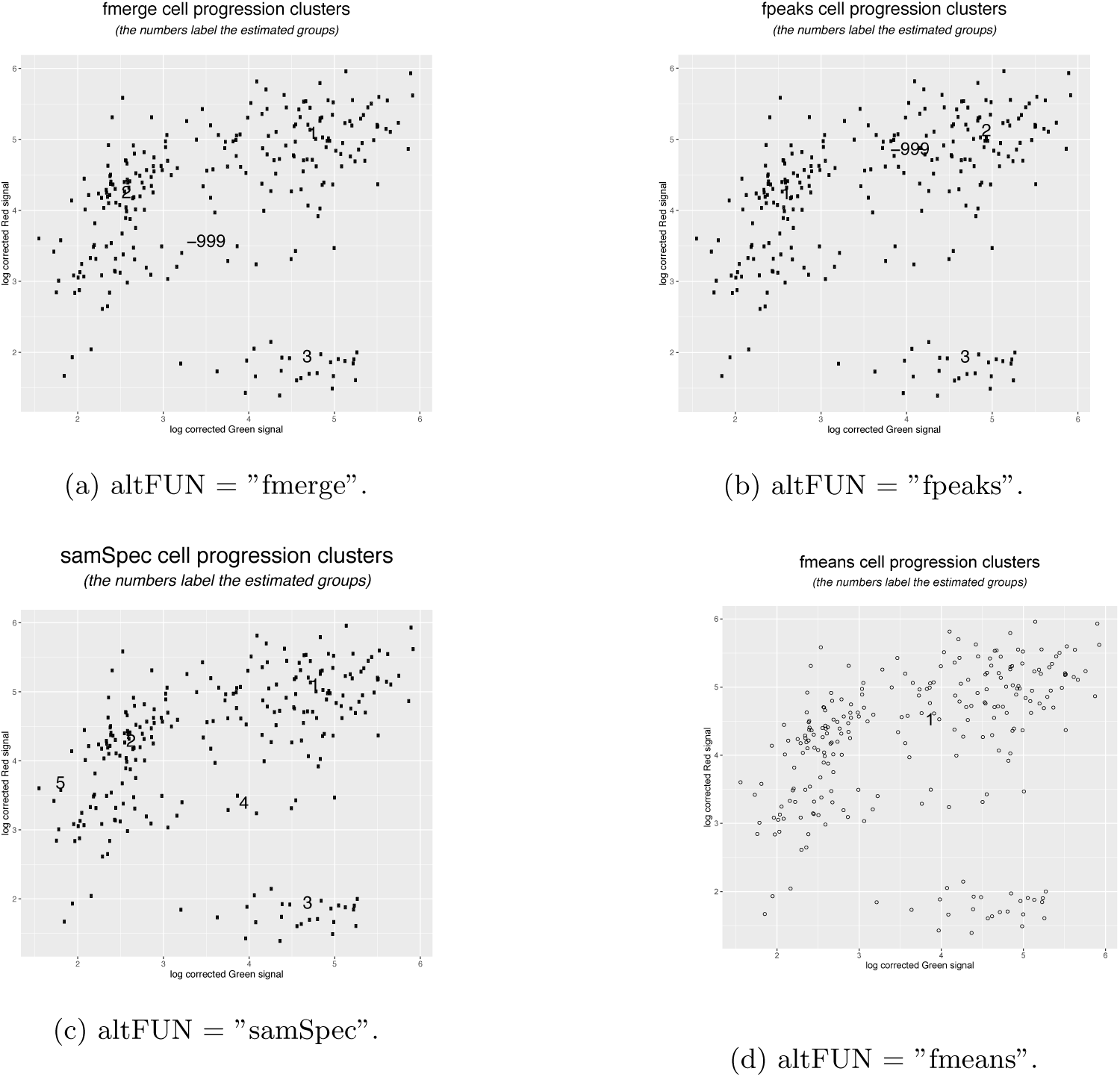
Estimated clusters by (a) **flowMerge**, (b) **flowPeaks**, (c) **SamSPECTRAL** and (d) **flowMeans**. All methods are implemented in **CONFESS**. The numbers indicate the different clusters.

## 6 Conclusions and further research

In this work we proposed a new direction to single-cell data analysis. It is based on the **CONFESS** statistical model that performs cell fluorescence signal estimation and modelling from images of living cells with fluorescent reporters determining the cell state. Via its main analytic steps of cell detection, signal correction and cell clustering / ordering, **CONFESS** enables the collection and accurate processing of cell morphology and cell signal information to further study the spatio-temporal dynamics of any system. The results can be directly integrated in the various transcriptome-driven statistical models to gain deeper understanding of the biological data in hand and improve the inference in gene regulation and transcriptional activation, coregulation networks, biomarker discovery and so on. Here, we showed the performance of our methodology on a large set of HeLa cells for which we were able to establish a clear cell progression path along the cell cycle. Our model can be used as a tool for benchmarking of single-cell RNA-seq based methods that perform cell ordering / state estimation from the transcriptome data only. In the future we intent to generalize our pipeline for other single-cell platforms (e.g. Dolomite; http://www.dolomite-microfluidics.com/), include 3-dimensional images for improved quality control and develop a consistent methodology that integrates the two sources of information, images and expression profiles, to obtain meaningful biological results on the cell cycle of HeLa and other cell types.

## Acknowledgements

We thank Dr Piero Carninci, Dr Charles Plessy and Mr Michael Böttcher for their support and helpful comments. This work was supported by the research grant from the Japanese Ministry of Education, Culture, Sports, Science and Technology (MEXT) to the RIKEN Center for Life Science Technologies.

## Authors contribution

EM and DHPL contributed equally in this work. EM conceived the idea, built the CONFESS model and wrote the manuscript. DHPL built **CONFESS** and **CONFESSdata** R packages for Bioconductor, helped in writing the manuscript and wrote the vignette. Correspondence should be addressed to EM.

## Supplementary Material

An .R file with the code that replicates the results of this work

